# Phosphorylation of AHR by PLK1 promotes metastasis of LUAD via DIO2-TH signaling

**DOI:** 10.1101/2023.07.31.551298

**Authors:** Chaohao Li, Derek B. Allison, Daheng He, Fengyi Mao, Xinyi Wang, Piotr Rychahou, Ibrahim A. Imam, Yifan Kong, Qiongsi Zhang, Yanquan Zhang, Jinghui Liu, Ruixin Wang, Xiongjian Rao, Sai Wu, Qing Shao, Chi Wang, Zhiguo Li, Xiaoqi Liu

## Abstract

Metastasis of Lung adenocarcinoma (LUAD) is a major cause of death in patients. Aryl hydrocarbon receptor (AHR) is an important transcription factor involved in the initiation and progression of lung cancer. Polo-like kinase 1 (PLK1), a serine/threonine kinase, is an oncogene that promotes the malignancy of multiple cancer types. Nonetheless, the interaction between these two factors and significance in lung cancer remains to be determined. Here, we demonstrate that PLK1 phosphorylates AHR at S489 in LUAD, which leads to epithelial-mesenchymal transition (EMT) and metastatic events. RNA-seq analyses show that type 2 deiodinase (DIO2) is responsible for EMT and enhanced metastatic potential. DIO2 converts tetraiodothyronine (T4) to triiodothyronine (T3), which then activates thyroid hormone signaling. In vitro and in vivo experiments demonstrate that treatment with T3 or T4 promotes the metastasis of LUAD, whereas depletion of DIO2 or deiodinase inhibitor disrupts this property. Taken together, our results identify the phosphorylation of AHR by PLK1 as a mechanism leading to the progression of LUAD and provide possible therapeutic interventions for this event.

## Introduction

Lung cancer is the leading cause of cancer-related deaths in the US and accounts for one-fifth of the total cancer mortalities among all populations, with approximately 350 deaths each day according to the latest cancer statistics [1]. The disease is preventable because most cases are related to cigarette smoking, which can be managed by smoking cessation and reduced exposure to environmental smoking [2]. However, considering the large number of patients and relatively low survival rate, lung cancer still places a heavy burden on society and the healthcare system. Among all forms of lung cancer, non-small-cell lung cancer (NSCLC) represents over 80% of all cases and can be further classified into three major subtypes: lung adenocarcinoma (LUAD), lung squamous cell carcinoma (LUSC), and large cell carcinoma [3]. Genetic mutations are the drivers of lung cancer, and drugs targeting these vulnerabilities are major treatments for patients with lung cancer. Although great achievements have been made in developing targeted therapies for lung cancer, current treatment options fail to control disease progression in the long run owing to the emergence of resistance [4]. Therefore, a deeper investigation of the etiology of lung cancer is necessary to better manage this disease.

Polo-like kinase 1 (PLK1) is a serine/threonine (S/T) protein kinase involved in the modulation of cell division at multiple phases, including mitotic entry, assembly of the kinetochore and spindle, centrosome maturation, activity of the anaphase-promoting complex, and cytokinesis [5–10]. Beyond its canonical role as a cell cycle regulator, PLK1 also possesses non-mitotic functions in other biological events, such as regulation of DNA damage repair, metabolism, and immune response [11–13]. Because of its wide functions, PLK1 is an oncogene frequently dysregulated in many cancers [14]. Specifically for NSCLC, PLK1 overexpression promotes lung cancer progression and is associated with poor patients’ outcomes [15, 16]. Considering its importance in lung cancer, novel therapies targeting PLK1 have been developed with observable efficacy in lung cancer cells and animal models [17, 18]. Although several PLK1 inhibitors have been tested in lung cancer patients, no effective PLK1 inhibitors are available for clinical cancer treatment, suggesting a more complicated and yet underexplored function in lung cancer.

The aryl hydrocarbon receptor (AHR) is a ligand-activated transcription factor and a well-known sensor for xenobiotics [19]. Originally discovered as the receptor for the environmental toxicants that results in the carcinogenesis of liver and skin cancer, it has been found that AHR is associated with multiple stages of tumorigenesis [20]. Upon binding to various ligands, AHR dimerizes with cofactors and translocates to the nucleus, where it initiates the transcription of downstream targets that are important for tumor initiation, proliferation, and metastasis. AHR has been shown to be overexpressed in many NSCLC cell lines and patients [21, 22]. In addition, high AHR activity is associated with poor overall survival and acquired resistance to treatment, suggesting its critical role as an oncogene in this deadly disease [23]. Therefore, AHR inhibitors are being developed and tested in clinical trials for the treatment of several solid tumors [24]. Although both PLK1 and AHR are closely related to lung cancer tumorigenesis, whether they cooperate to accelerate tumor progression remains unknown. Here, we show that AHR is a direct substrate of PLK1 and that phosphorylation of AHR in LUAD promotes epithelial–mesenchymal transition (EMT) to enhance metastatic potential. Mechanistically, AHR phosphorylation elevates the expression of type 2 deiodinase (DIO2) and activates thyroid hormone (TH) signaling. Overall, our findings reveal a novel mechanism linking two important tumor promoters in lung cancer and provide translational value for cancer interventions.

## Methods

### In vitro kinase assay

Human AHR CDS was cloned into the pGEX-KG vector. Protein purification of fragmented AHR was performed using the BL21 system and Glutathione Sepharose 4B (Cytiva, 17075601). Full-length human AHR was purified using Pierce Anti-HA Magnetic Beads (Thermo, 88836) from 293T cell line transfected with HA-AHR plasmids. The detailed conditions of the in vitro kinase assay have been previously described [25]. Briefly, the reaction mixture was prepared with substrates, reaction buffers, and radioactive ATP and then incubated at 30°C for 30 mins, followed by SDS-PAGE gel electrophoresis. The gel was stained with 0.1% Coomassie Blue, destained, dried under vacuum, and subjected to autoradiography. Validation of antibodies by fragmented AHR was performed using nonradioactive ATP, and 0.5% Ponceau S was used for gel staining. After destaining with 1x TBST buffer, the gel was subjected to immunoblotting.

### Prediction of phosphorylation and mutagenesis of plasmids

Prediction of phosphorylation sites by PLK1 was performed with GPS 5.0 and GPS-Polo 1.0 [26, 27]. Mutagenesis of the HA-AHR plasmid was performed using Q5 Site-Directed Mutagenesis Kit (New England Biolabs, E0554S).

### Cell culture

The original A549 cell line was kindly provided by Dr. Qiou Wei at the University of Kentucky, and other cell lines were purchased from ATCC. Generation of V, WT, SA, and SD cell lines stably expressing different mutations of AHR was established by depleting endogenous AHR, then overexpressing HA-AHR plasmids or their empty vector. 200ug/ml hygromycin was used to maintain cells. TET-PLK1 and TET-shPLK1 cell lines were generated using the TET-FLAG-PLK1 and TET-shPLK1 plasmids, respectively. Puromycin (2 ug/ml) was used to maintain cells. SD-shDIO2 cells were generated using shRNA plasmids. Puromycin (2 ug/ml) was used to maintain cells. A549 cells expressing firefly luciferases and GFP were generated using pCDH- EF1-Luc2-cG-BSD. The A549 and H1299 cell lines and their derivatives were cultured in RPMI 1640 medium. 293T cells were cultured in high-glucose DMEM. The culture conditions for all the cell lines were 10% FBS, 5% penicillin-streptomycin, 37 °C, and 5% CO2. All the cell lines were within 50 passages and tested negative for mycoplasma contamination.

### Wound healing assay

Cells were seeded in 6-well plates and cultured in complete medium until 100% confluence. 200ul tips were used to scratch the middle areas of the wells. Images were taken at three independent areas every 24 h with the Nikon microscope and analyzed using Fiji [28]. Before taking pictures, the cells were washed with 1x PBS twice to remove floating cells. Cells were cultured in a medium with the indicated chemicals, and the concentration of FBS was 0.5%, which was optimized to avoid proliferation. All experiments were repeated three times and one was shown here.

### Transwell assay

#### Migration assay

Cells (5 × 10^4^) were seeded on top of 8.0 µm 24-well inserts (Corning, 353097) with 100ul medium containing 0.5% FBS and the indicated chemicals. 600ul complete medium were added to the bottom of each well to induce chemotaxis. After 24 hours, cells were fixed with 70% ethanol for 30 mins and stained with 0.1% crystal violet for 30 mins, then washed with tap water until clear. Cells on top of the inserts were removed using cotton swabs. Images were taken with the Nikon microscope and analyzed using Fiji. **Invasion assay.** Matrigel (Corning, 354234) was diluted with medium containing 0.5% FBS at a ratio of 1:50. 50ul diluted mixture was used to coat the top of the inserts at 37 °C for 30 mins. Subsequent procedures were the same as those used for the migration assay. All experiments were repeated three times and one was shown.

### 2D growth assay

Cells (2 × 10^3^ cells/well in 200µl) were seeded in 96-well plates. The medium was refreshed with the indicated concentrations of the chemicals every 72 h. On the indicated day, AquaBluer (MultiTarget Pharmaceuticals LLC, 6015) reagent was mixed with the medium at a ratio of 1:100, the medium was aspirated and 100ul mixture was added to each well, followed by incubation for 4 h. Fluorescence intensity at 540ex/590em was read by the GloMax Discover plate reader (Promega). All experiments were repeated three times and one was shown.

### 3D culture

#### 3D invasion assay

Embedded 3D cultures were used in the assay. Briefly, Matrigel was diluted with medium containing 2% FBS at a ratio of 1:1, then 100ul diluted Matrigel was used to coat 24-well plates at 37 °C for 30 mins. Next, the cell suspension (5 × 10^3^ cells/well) was mixed with diluted Matrigel at a ratio of 1:10, then 300ul mixture was added to the coated 24-well plates and incubated at 37 °C for 30 mins. Finally, 500ul complete medium with the indicated chemicals was added to each well, and the culture was maintained for 6 to 8 days. The medium and chemicals were refreshed every two days. Images were taken with the Nikon microscope and analyzed using Fiji. **3D spheroid formation and growth assay.** The assay has been previously described with slight modifications [29]. Briefly, cells (1 × 10^3^ cells/well in 100µl) were diluted in 80% complete medium with 20% Matrigel, seeded onto 96-well plates and incubated at 37 °C for 30 mins. Following Matrigel solidification, 50ul complete medium with indicated chemicals was added to each well and kept culture for 6 to 8 days. The medium and chemicals were refreshed every two days. On the last day, spheres were imaged with the Nikon microscope and analyzed using Fiji. Alternatively, AquaBluer reagent was mixed with the medium at a ratio of 1:100, the medium was aspirated and 50ul mixture was added to each well, followed by incubation for 4 h. The fluorescence intensity at 540ex/590em was read by the GloMax Discover plate reader.

### Animal experiments

All animal experiments used in this study were approved by the University of Kentucky Division of Laboratory Animal Resources. 6-week-old Athymic nude mice (The Jackson Lab, 002019) were used for all experiments. All tissues were formalin-fixed and paraffin-embedded (FFPE). Whole slides were scanned using the Aperio Digital Pathology Slide Scanner (Leica). Images were viewed and quantified using ImageScope (Leica). **Intravenous injection of V, WT, SA, and SD cells.** Cells (5 × 10^5^ in 100µl PBS) were intravenously injected into each nude mouse. All mice were euthanized at week 12 based on a previously reported study [30]. FFPE tissues were sent to the histological laboratory for hematoxylin and eosin (H&E) staining. **Intravenous injection of SD and SD-shDIO2 cells.** Iopanoic acid (IOP) was dissolved in 10% DMSO and 90% corn oil. Cells (5 × 10^5^ in 100µl PBS) were intravenously injected into each nude mouse. Mice in the IOP treatment group were injected with SD cells and treated with 25 mg/kg IOP daily via oral gavage. In vivo imaging of the lung metastasis was performed at week 11. All mice were euthanized at week 12 and FFPE tissues were sent to the histology laboratory for H&E staining. **Subcutaneous injections of V, WT, SA, and SD cells.** Cells (2 × 10^6^ in 100µl PBS) were subcutaneously inoculated into each mouse. The tumor sizes were measured weekly using a digital caliper. Tumor volumes were calculated using the formula: V= L × W^2^ × 0.52, where V is volume (mm^3^), L is length (mm), and W is width (mm). The treatment was stopped when the tumor volume reached 1000mm^3^. At least 100ul whole blood was collected from the heart of each mouse upon euthanasia and analyzed for GFP^+^ tumor cells.

### Flow cytometry

All data were acquired on the BD Symphony A3 analyzer (BD Biosciences) and analyzed using FlowJo software. **Cell cycle analysis.** The cell cycle was analyzed using the Vybrant DyeCycle Violet Stain (Thermo, V35003). Briefly, Cells (1 × 10^6^) in 1ml complete media were stained with 1ul dye at 37°C for 30 mins. The fluorescence intensity was acquired at 405ex/440em**. Analysis of GFP^+^ tumor cells.** Blood was collected via terminal cardiac puncture of the right ventricle using a 23-G needle attached to a 3-mL blood collection tube precoated with EDTA. Samples were subjected to 1X RBC Lysis Buffer (BioLegend, 420301), washed with PBS, and resuspended in 400ul PBS. Circulating tumor cells were collected at 488ex/510em.

### In vivo imaging of lung metastasis

Mice were intraperitoneally injected with 150 mg/kg D-Luciferin (GoldBio, LUCK). 10 mins later, mice were anesthetized in chamber filled with isoflurane and subject to imaging on Lago X imaging system at 570nm wavelength. The results were analyzed using Aura Imaging Software.

### Statistical analysis

All results were analyzed with the statistical functions in GraphPad Prism 8, except the results in S5D, which was analyzed by the package "lme4" in R-4.2.2. Normality and variance of results were checked, and appropriate statistical analyses were performed. Statistical significance was set at p values < 0.05. The detailed methods can be found in the corresponding figure legends.

## Results

### PLK1 phosphorylates AHR at S489

Preliminary screening of human AHR sequences using software indicated a phosphorylation event by PLK1. To investigate this possibility, we cloned the full-length human AHR cDNA sequence into four fragments (F1-F4) based on its functional domains (Fig. 1A), and radioactive phosphorus- 32 (^32^P) kinase assays were performed. The results showed that only F3 had a ^32^P signal, which disappeared upon treatment with the PLK1 inhibitor onvansertib (ONV), suggesting that phosphorylation of AHR occurred in F3 (Fig. 1B). A visual examination of F3 revealed that the sequence around S489 was highly congruent with the previously identified consensus phosphorylation motif of PLK1 [31] (Fig. 1C). Together with the potential sites predicted by the software (Fig. 1D), we mutated each individual S or T to A in F3, and then performed ^32^P kinase assays to determine which mutation could lead to reduction of the ^32^P signal. Mutation of S489 to A (SA) resulted in the greatest reduction in the ^32^P signal (Fig. 1E). Additional ^32^P kinase assays with triplicate F3 or full-length human AHR purified from 293T cells confirmed SA as a bonafide phosphorylation site (Fig. 1F). Co-immunoprecipitation of PLK1 and AHR in 293T cells revealed an interaction between the two proteins (Fig. 1G), further confirming the kinase-substrate relationship. Notably, this site was conserved among higher animals, but not between humans and mice, suggesting that SA was a unique PLK1 phosphorylation site in humans (Fig. 1H). Based on these results, we argued that human AHR was a substrate of PLK1.

**Figure 1.**
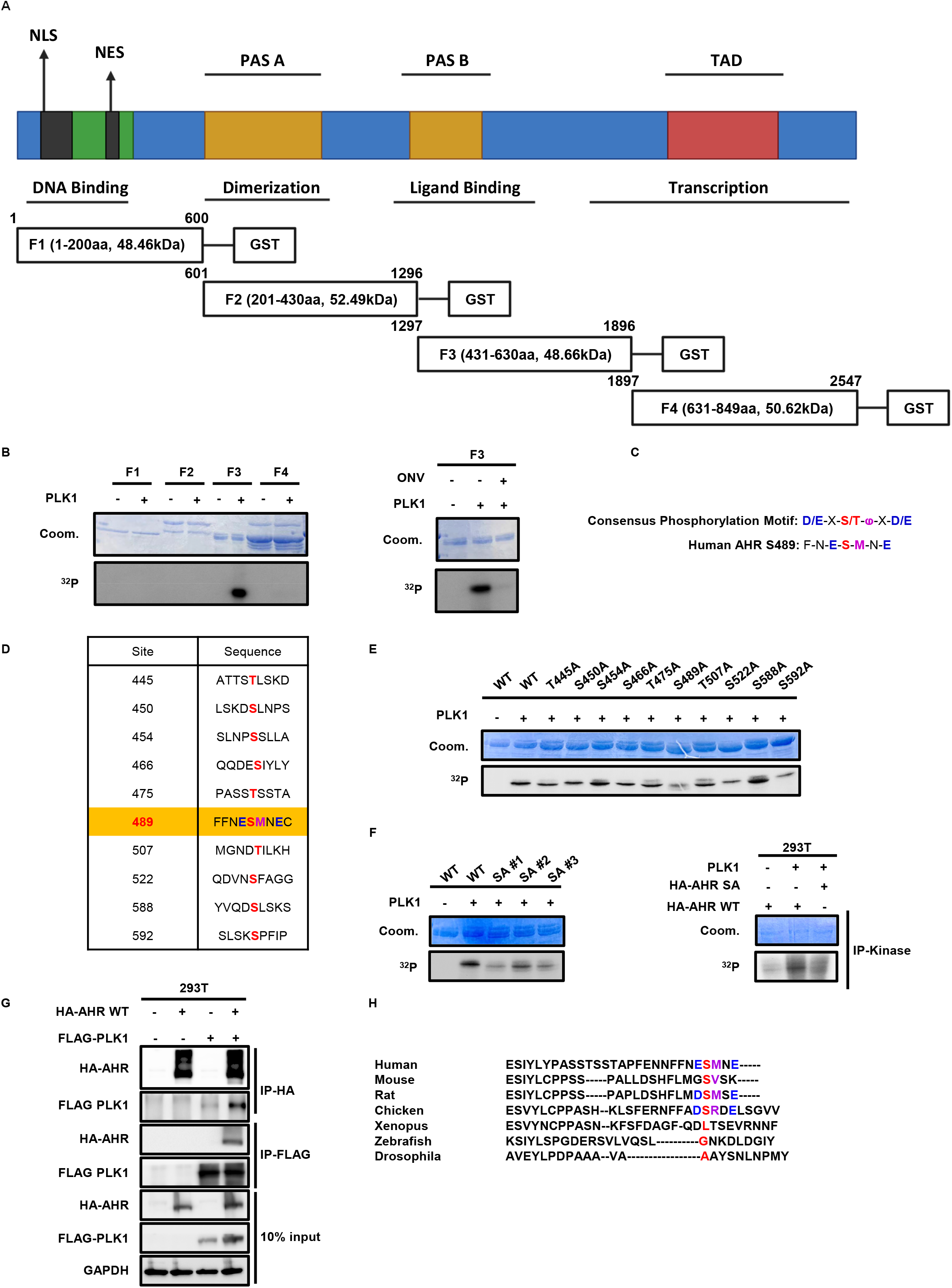
In vitro kinase assay identifies AHR as the PLK1 substrate. **A**, Fragmentation of full-length AHR into F1, F2, F3, and F4 based on its functional domains. The C-termini of each fragment is tagged with glutathione s-transferase (GST). NLS, nuclear localization signal. NES, nuclear export signal. PAS, Per-Arnt-Sim. TAD, transactivation domain. **B,** In vitro kinase assay with four fragments. Coom, Coomassie Blue. ^32^P, radioactive phosphorus-32. ONV, onvansertib. **C,** Previous identified consensus phosphorylation motif of PLK1 substrates and S489 site of F3. **D,** Potential PLK1 phosphorylation sites identified using the software. **E,** In vitro kinase assay with wild-type (WT) or mutated F3. **F,** In vitro kinase assay with F3 or HA- tagged full-length AHR (HA-AHR) with the S489A (SA) mutation. **G,** Co-immunoprecipitation of HA-AHR and FLAG-tagged PLK1 (FLAG-PLK1) in 293T cells. **H,** Alignment of consensus sequences around S489 in different species.

### AHR phosphorylation is associated with LUAD progression

To confirm this phosphorylation event in cells, we generated homemade antibodies targeting the phospho-S489 epitope of AHR (p-AHR) and verified their efficacy using purified AHR or in cell models (Fig. 2A). The results showed that the two antibodies detected p-AHR in purified F3 or in 293T cells transfected with full-length proteins (Fig. 2B), confirming their specificity and phosphorylation in cells. Hence, we used an antibody with a lower background signal (#1) in our experiments. Next, we detected p-AHR in different subtypes of human lung cell lines (Fig. 2C). While AHR in normal lung cell lines was weakly phosphorylated, cancer cell lines generally displayed stronger p-AHR. Compared to normal lung cells, LUAD cell lines showed consistent overexpression of both PLK1 and AHR, which was not observed in LUSC or small cell lung cancer cell lines. Furthermore, survival analysis of the TCGA-LUAD dataset revealed that patients with high PLK1 and high AHR had the worst outcomes regarding their four different survival results (Fig. 2D, Table. S1). In contrast, this phenomenon was not observed in the TCGA-LUSC dataset (Fig. S1, Table. S2), suggesting the critical clinical impact of this phosphorylation event in LUAD. Accordingly, we focused on LUAD and used the A549 cell line, which displayed the highest p- AHR level, to dissect the functional significance of phosphorylation. Treatment of the A549 cell line with nocodazole (NOC) elevated PLK1 and increased p-AHR, which was reversed by ONV treatment, and NOC release experiment further supporting this regulation, in which the decrease of PLK1 was accompanied by the decline of p-AHR (Fig. 2E). To provide further evidence, we established tetracycline-controlled overexpression of PLK1 (TET-PLK1) and depletion of PLK1 (TET-shPLK1) systems in the A549 cell line. Treatment with doxycycline to overexpress or deplete PLK1 witnessed an elevation or reduction of p-AHR (Fig. 2F), confirming the phosphorylation of AHR by PLK1. Taken together, these data demonstrated that phosphorylation might be associated with disease outcomes of LUAD.

**Figure 2.**
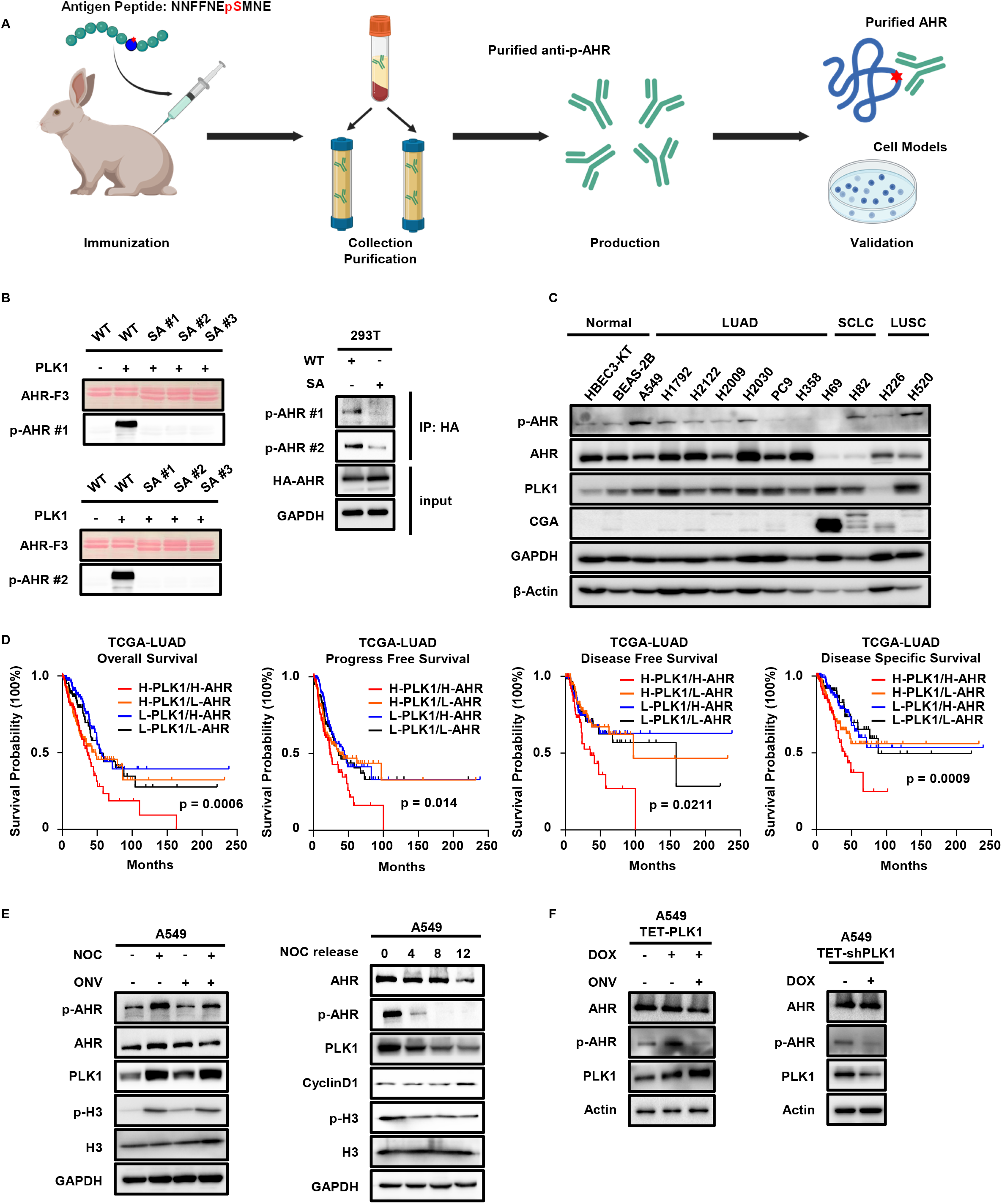
AHR phosphorylation is implicated in LUAD progression. **A**, Illustration of homemade anti-p-AHR antibodies and validating experiments. **B,** Validation of specificities of two anti-p-AHR antibodies with F3 or HA-AHR. **C,** Immunoblotting (IB) to detect p-AHR in normal lung cells and different subtypes of lung cancers. LUAD, lung adenocarcinoma. LUSC, lung squamous cell carcinoma. SCLC, small cell lung cancer. CGA, Chromogranin A, marker of SCLC. **D,** Kaplan-Meier survival curves of TCGA-LUAD patients. Based on the median expressions of PLK1 and AHR, patients are separated into four groups: High PLK1/High AHR (H-PLK1/H-AHR, PLK1/AHR > median, n = 102/102/95/53 from left to right), High PLK1/Low AHR (H-PLK1/L-AHR, PLK1 > median/AHR < median, n = 146/146/132/80 from left to right), Low PLK1/High AHR (L-PLK1/H-AHR, PLK1 < median/AHR > median, n = 147/147/142/101 from left to right), Low PLK1/Low AHR (L-PLK1/L-AHR, PLK1/AHR < median, n = 106/106/97/63 from left to right). Statistical method: Log-rank test. **E,** IB to detect p-AHR in nocodazole (NOC) treatment and release experiments in A549 cells. **F,** IB against p-AHR in A549 cells with tetracycline-controlled induction of FLAG-PLK1 (TET-PLK1) or depletion of PLK1 (TET-shPLK1). DOX, doxycycline.

### Phosphorylation of AHR stimulates the EMT and metastatic potential of LUAD

To investigate the biological significance of p-AHR in LUAD, we examined several hallmarks of cancer in LUAD (Fig. 3A). By introducing an empty vector (V), wild-type AHR (WT), SA, or phosphomimetic AHR with a mutation of S489 to D (SD) in A549 cells (Fig. 3B), we established stable cell lines and used them to study the function of phosphorylation. We found that WT and SD cells had stronger migratory abilities as shown by the wound healing assay, with SD cells displaying the strongest capacity (Figs. 3C-3D), suggesting that phosphorylation might enhance the metastatic potential of LUAD. Of note, PLK1 was previously reported to be a promoter of lung metastasis [15, 32], and this was supported by our wound healing assays with TET-PLK1 and TET-shPLK1 cells (Fig. S2A), in which overexpression or depletion led to the accelerated or decelerated closure of scratches, respectively. In addition, high AHR was also shown to be related to disease recurrence and distant metastasis in patients with LUAD [23]. Based on these preliminary results, we hypothesized that PLK1 phosphorylated AHR to promote the invasive properties of LUAD. To test this hypothesis, we performed transwell migration and invasion assays. Besides, 3D invasion assay, which cultured cells in matrix-embedded medium, was also applied to provide a more realistic condition to mimic the in vivo environment, in which tumor cells needed to cross the barrier of the extracellular matrix before the initiation of distant metastasis. Compared to V and SA cells, WT and SD cells had stronger abilities to penetrate the transwell membranes and the surrounding matrix, with SD cells consistently showing the most significant difference (Fig. 3E), supporting the notion that phosphorylation was a prerequisite for promoting metastasis. To rule out the possibility that the observed phenotypes were due to the distinct proliferation rates, we checked the growth properties of those cells. While 2D growth assay showed statistical significance between several groups, the difference was very minimal (less than 10%) to be considered as important (Fig. 3F). Additional experiments, including 3D spheroid formation, 3D growth assay, and cell cycle analysis, also failed to detect large differences (Figs. 3G, S2B-S2C). Apparently, the enhanced metastatic potential was not likely due to the differences in growth properties. These data were recapitulated in another LUAD cell line, H1299 (Figs. S3A- S3G), further confirming the reliability of this phenomenon. Notably, PLK1 is an established inducer of EMT in multiple cancers [33], which is a precondition for metastatic events. To test whether this was responsible for the effect of phosphorylation, we detected epithelial markers (E- Markers) and mesenchymal markers (M-Markers) in these cells. We found that WT and SD cells expressed higher M-Markers of N-Cadherin (N-Cad) and Vimentin (VIM), but lower E-Markers of E-Cadherin (E-Cad), with SD cells expressing highest M-Markers and lowest E-Markers (Fig. 3H), confirming the contribution of phosphorylation to promote EMT. Taken together, these data supported the hypothesis that PLK1 phosphorylated AHR to induce EMT and enhance the metastatic potential of LUAD cells.

**Figure 3.**
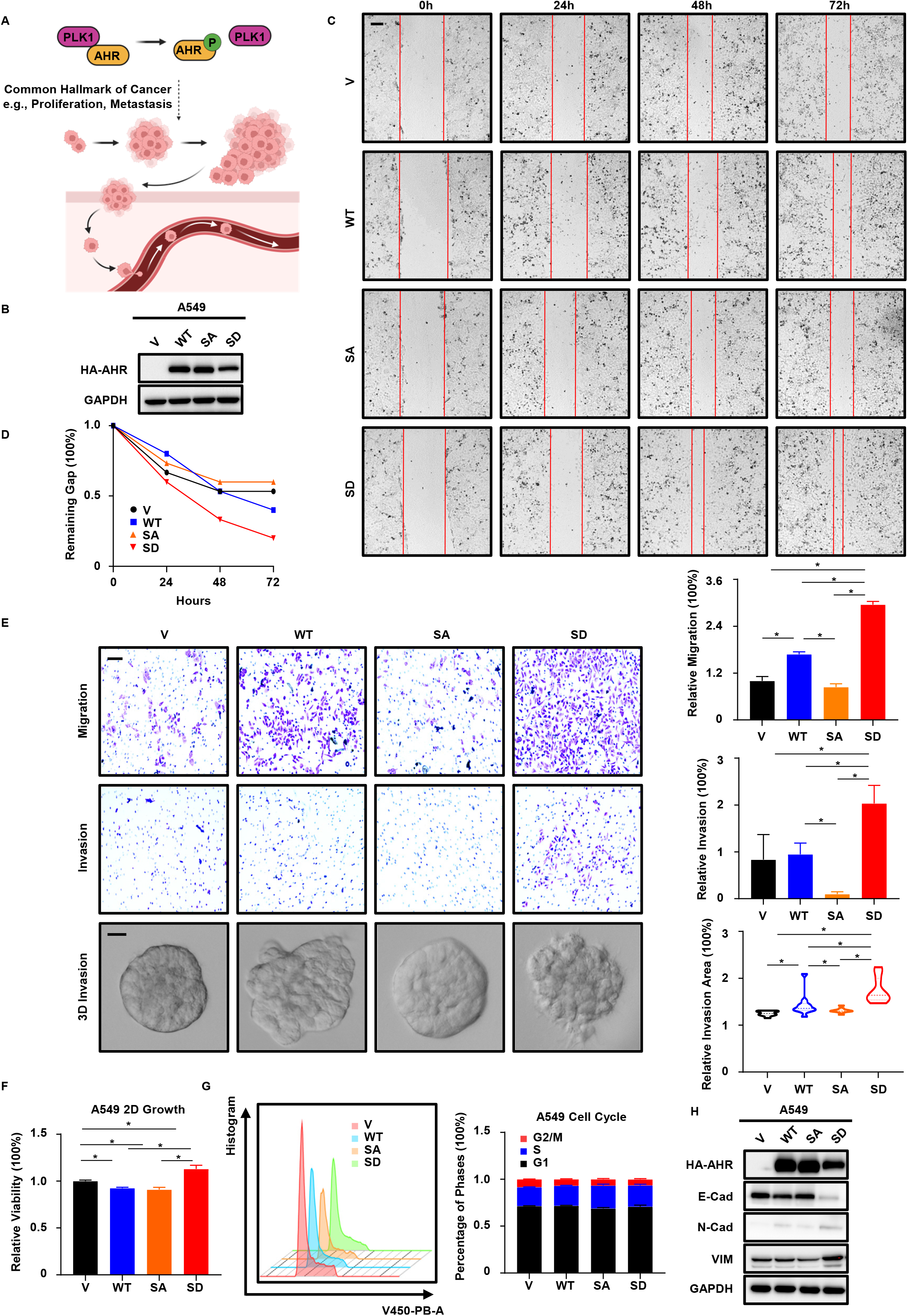
Phosphorylation of AHR promotes EMT and metastasis of LUAD. **A**, Possible outcomes of phosphorylation of AHR by PLK1. **B,** IB to verify establishment of A549 cells stably expressing empty vector (V), WT HA-AHR and HA-AHR with SA or S489D (SD) mutations. **C, D,** Wound healing assay with V, WT, SA, SD cells. Results are normalized to 0h. Scale bar, 250µm. **E,** Transwell migration, invasion and 3D invasion assays with V, WT, SA, SD cells. Results are normalized to V and shown as mean ± SD (n = 3 for transwell, n = 10 for 3D invasion). Scale bar (upper two panels), 100µm. Scale bar (bottom panel), 20µm. Statistical methods: two-tailed unpaired Welch’s t test (invasion assay, SD-SA); two-tailed unpaired t test (remaining comparisons in transwell assays); two-tailed Mann-Whitney test (3D invasion assay). **F,** 3-day 2D growth assay with V, WT, SA, SD cells. Results are normalized to day 0 and shown as mean ± SD (n = 8). Statistical method: Welch ANOVA test following multiple comparisons. **G,** Cell cycle analysis of V, WT, SA, SD cells by flow cytometry. Results of each phase are scaled to 100% and shown as mean ± SD (n = 4). **H,** IB to detect epithelial–mesenchymal transition (EMT) and the associated epithelial markers (E-Markers), mesenchymal markers (M-Markers) in V, WT, SA, SD cells. **\***, p < 0.05.

### AHR phosphorylation promotes LUAD metastasis in vivo

Metastasis of primary tumors to distant tissues involves several critical steps, including local invasion, intravasation, survival in the circulating system, extravasation, and residency at secondary sites [34]. To mimic this process and support our in vivo findings, we designed two animal experiments to assess the metastatic abilities of these cells (Fig. 4A). First, we introduced GFP into the stable cell lines and subcutaneously inoculated cells into nude mice, which mimicked the first two steps of metastasis. Tumor growth over 2 months was similar among the four groups, and the final tumor weights displayed a slight difference between the V and SA cells (Figs. 4B- 4C), indicating minimal differences in the cell growth properties in vivo. However, analysis of blood samples showed that mice inoculated with SD cells had a higher percentage of GFP-positive cells compared to other groups (Figs. 4D, S4), suggesting a stronger tendency of SD cells to invade and intravasate into circulating systems. Next, we intravenously injected four cell lines into nude mice and monitored the extent of lung residency, which mimicked the last three steps of metastasis. In keeping with previous results, WT and SD cells showed more metastasis to lung tissues compared to V and SA cells, with the SD group exhibiting the largest metastatic lesions (Fig. E). In summary, these data demonstrated that phosphorylation stimulated metastasis of LUAD in vivo.

**Figure 4.**
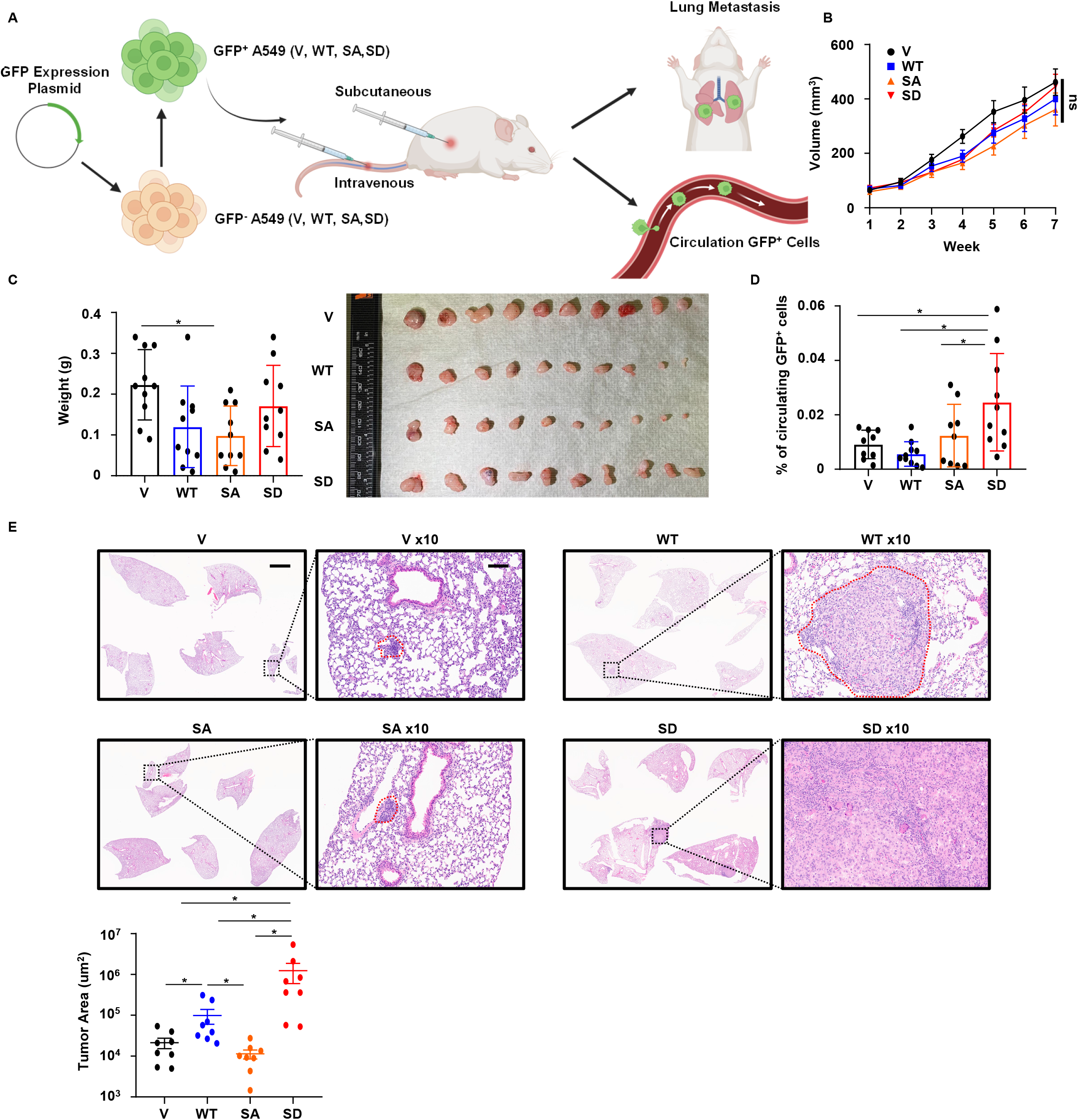
AHR phosphorylation promotes LUAD metastasis in vivo. **A**, Schematic representation of in vivo metastasis experiments. GFP^+^, GFP positive. GFP^-^, GFP negative **B,** Tumor growth of V, WT, SA, SD cells over 7 weeks. Results are shown as mean ± SEM (n = 10). Statistical method: one-way ANOVA test. **C,** Final tumor weights and photograph of V, WT, SA, SD tumors. Results are shown as mean ± SD (n = 10). Statistical method, one-way ANOVA following multiple comparisons. **D,** Flow cytometry analysis of circulating GFP^+^ tumor cells. Results are shown as mean ± SD (n = 10 for WT, SD and n = 9 for V, SA). Statistical method: one-tailed unpaired Welch’s t test. **E,** Hematoxylin and eosin (H&E) staining of lung tissue sections of mice intravenously injected with V, WT, SA, SD cells. Scale bar (origin), 2mm. Scale bar (10x), 80µm. Results of quantification are shown as mean ± SEM (n = 8). Statistical method: two-tailed Mann-Whitney test. **\***, p < 0.05. ns, not significant.

### Sequencing analysis identifies DIO2 as a molecular determinant of metastasis in LUAD

To investigate the mechanisms underlying phosphorylation and metastasis, we performed RNA-seq using these cell lines (Fig. 5A). Gene Set Enrichment Analysis (GSEA) of hallmark gene sets by differentially expressed genes (DEGs) showed that EMT was among the top-upregulated pathways in SD cells compared to WT and SA cells (Figs. 5B, S5A-S5B), consistent with previous findings. To further explore the molecular determinant and match with results in cellular experiments, we applied a filter so that only genes expressing highest in SD cells but lowest in SA cells were selected (SD > WT > SA). We then ranked these genes according to their significance (p values), and a list of top upregulated genes was shown together with their correlations with PLK1 in the TCGA-LUAD dataset (Fig. 5C). RNA-seq results indicated two targets that were significantly upregulated in SD cells (q values < 0.05, marked with asterisks), but only type 2 deiodinase (DIO2) was consistent in TCGA-LUAD and positively correlated with PLK1 (Spearman p values < 0.05, marked with asterisk and red color). To further explore the significance of DIO2, we analyzed its relationship with PLK1 and disease outcomes in TCGA-LUAD. DIO2 was slightly correlated with PLK1 in normal samples but became more closely in tumor samples (Fig. S5C). Compared to normal tissues, both PLK1 and DIO2 levels were elevated in tumor cells and at higher stages of disease (Fig. S5D). Of note, DIO2 was only higher in LUAD of IIB stage, when local metastasis started to appear, suggesting its importance during the early onset of metastasis. To investigate the correlation between DIO2 expression and survival, we separated patients into low DIO2 (L-DIO2), intermediate DIO2, and high DIO2 (H-DIO2) groups. Compared with the intermediate and low DIO2 groups, the H-DIO2 group had a higher rate of metastatic events (Fig. 5D). In addition, patients in L-DIO2 group had better survival outcomes compared to those in H-DIO2 group, with strong improvement of overall survival and disease free survival (Figs. 5E, S5E-S5F, Table. S3). Conversely, DIO2 was not positively correlated with PLK1 in TCGA-LUSC, and patients in the H-DIO2 and L-DIO2 groups had indistinguishable differences in survival outcomes (Figs. S5G-S5I, Table. S3). Furthermore, H-DIO2 closely correlated with higher M-Markers and lower E-Markers (Figs. 5F, S6), consistent with previous results that SD cells displayed a mesenchymal-like state favoring the initiation of metastasis. Indeed, DIO2 was recently shown to promote the progression and invasiveness of skin cancer by inducing EMT [35]. However, this was not determined in patients with LUAD. Based on these preconditions, we argued that DIO2 was critical for the metastatic phenotype of SD cells. We verified the overexpression of DIO2 by detecting the RNA and protein levels in the four cell lines (Fig. 5G). Besides, modulation of PLK1 in TET-FLAG and TET-shPLK1 cells confirmed the regulation of DIO2 by PLK1 (Fig. 5H). Taken together, our results suggested that elevation of DIO2 upon phosphorylation was responsible for enhanced metastatic abilities in LUAD.

**Figure 5.**
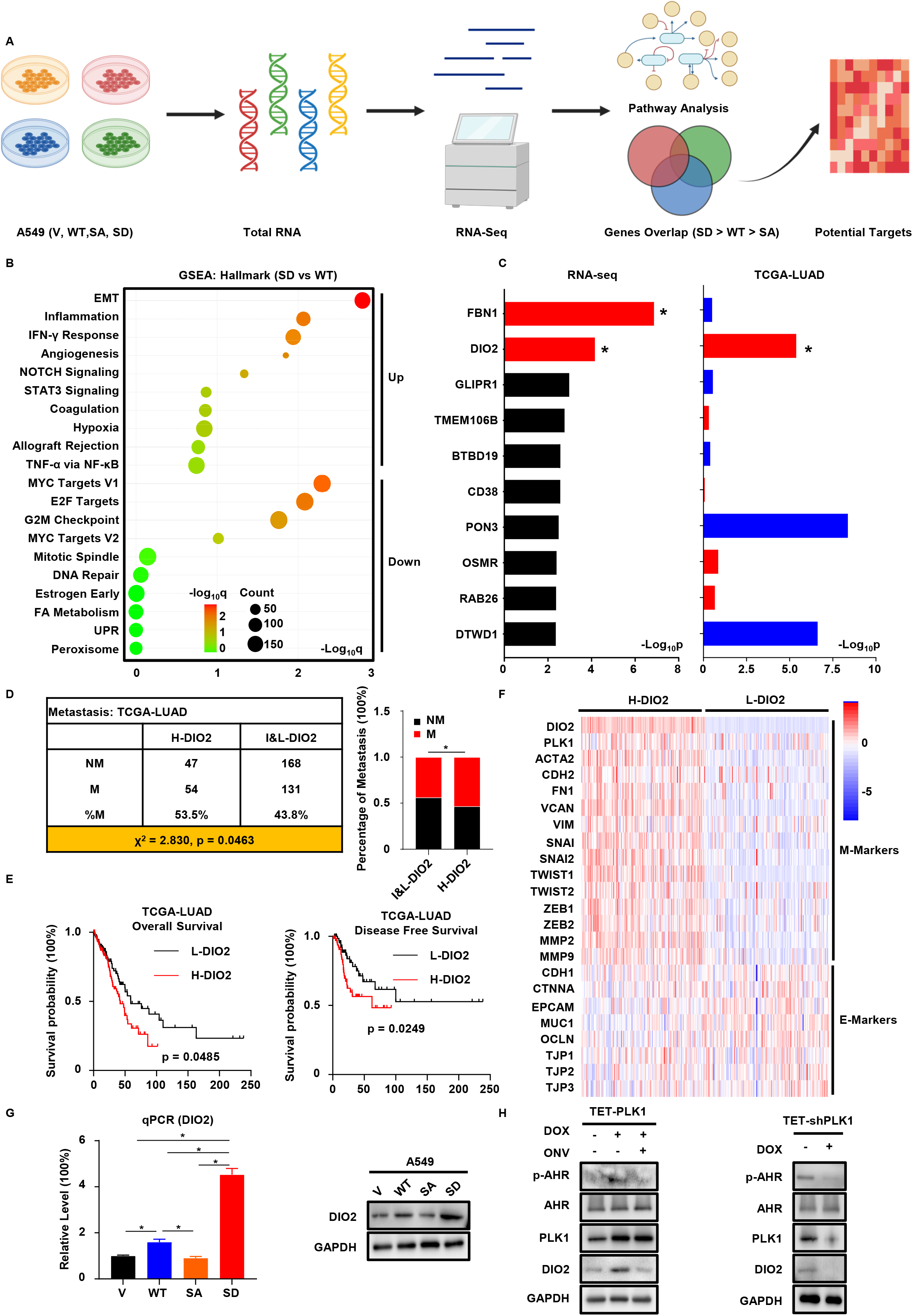
RNA-seq indicates DIO2 is responsible for metastasis. **A**, Simplified graphic representation of RNA-seq analysis. **B,** Bubble plot (SD vs WT) of top upregulated and downregulated pathways in SD cells, identified by Gene Set Enrichment Analysis (GSEA) with hallmark gene set. Log_10_q, log_10_-transformed false discovery rate values (q). **C,** Barplots of top upregulated genes (SD > WT > SA) identified from RNA-seq and their spearman correlations with PLK1 from TCGA-LUAD. Log_10_p, log_10_-transformed p values. Genes with q values < 0.05 in RNA-seq are marked with asterisks and red colors. Genes with positive correlations and p values < 0.05 in TCGA-LUAD are marked with asterisks and red color. Negative correlations in TCGA-LUAD are marked with blue color, regardless of the significance. **D,** Analysis of metastatic events of TCGA-LUAD patients. Based on the 3^rd^ quartile expression of DIO2, patients are separated into two groups: High DIO2 (H-DIO2, >= 3^rd^ quartile, n = 101), Intermediate/Low DIO2 (I&L-DIO2, < 3^rd^ quartile, n = 299). Metastatic events are characterized by N > 0 or M1. NM, non-metastatic. M, metastatic. Statistical method: one-sided Chi-square test. **E,** Kaplan-Meier survival curves of patients in E. Only patients with survival information are plotted. H-DIO2, n = 125/67 from left to right. L-DIO2, n = 125/69 from left to right. Statistical method: Log-rank test. **F,** Heatmap to show the expression of EMT markers in TCGA-LUAD patients. Based on the 1^st^ and 3^rd^ quartile expression of DIO2, patients are separated into two groups: H-DIO2 (n = 128) and Low DIO2 (L-DIO2, <= 1^st^ quartile, n = 128). **G,** qPCR and IB to detect the expression of DIO2 in V, WT, SA, SD cells. Results of qPCR are normalized to V and shown as mean ± SD (n = 8). Statistical methods: two-tailed unpaired t test (WT-SA, WT-SD); two-tailed unpaired Welch’s t test (WT-V, SD-V, SD-SA). **H,** IB to detect DIO2 in A549 cells with TET-PLK1 and TET-shPLK1. **\***, p < 0.05.

**Figure 6.**
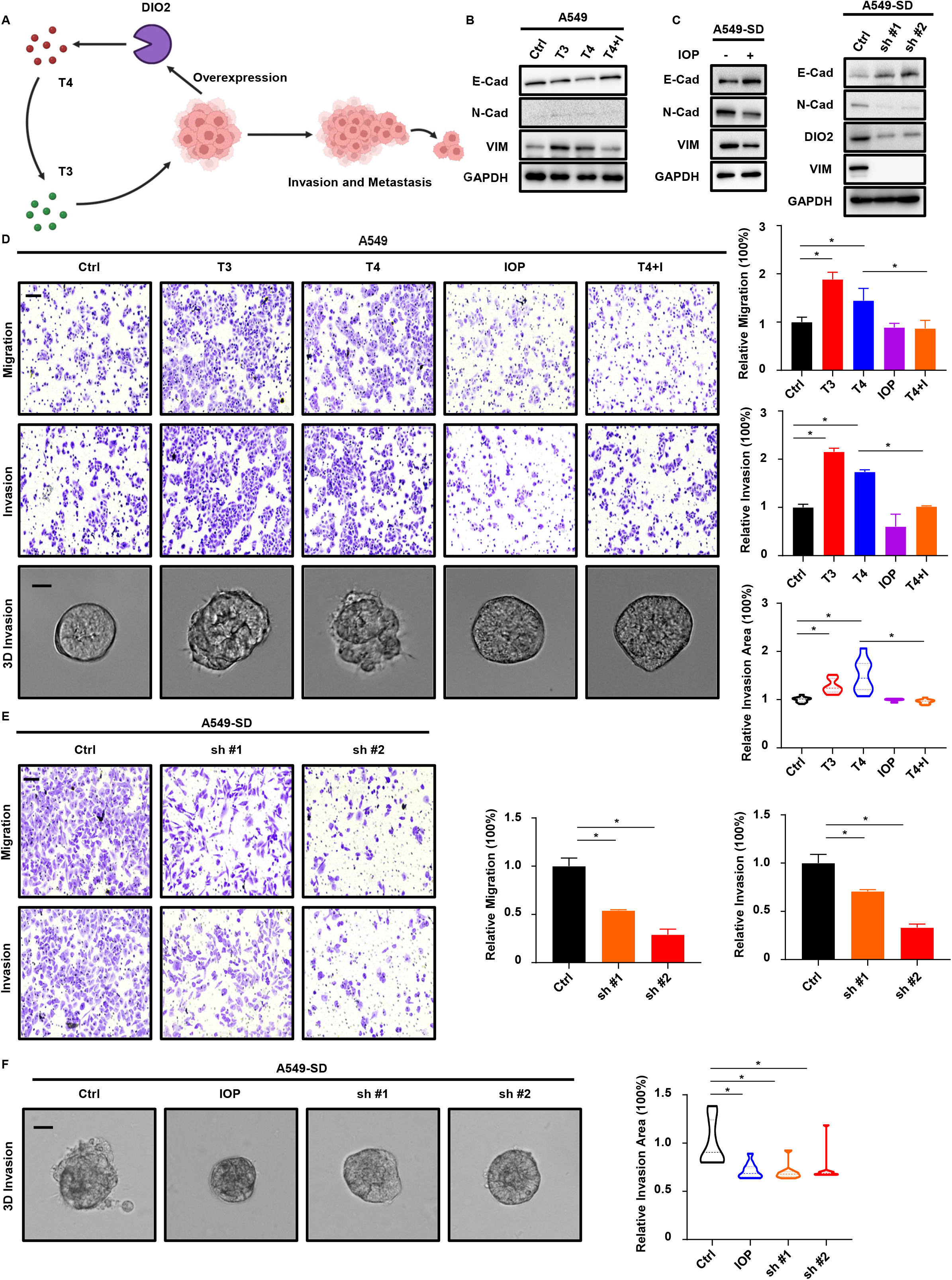
DIO2 and TH promote metastasis of LUAD. **A**, Function of DIO2 in thyroid hormone (TH) metabolism, represented by graph. T4, tetraiodothyronine. T3, triiodothyronine. **B,** IB to detect EMT markers in A549 cells treated with T3, T4 or T4 plus DIO2 inhibitor iopanoic acid (T4+I). Cells treated with DIO2 serve as control (Ctrl). **C,** IB to detect EMT markers in SD cells treated with DIO2 inhibitor iopanoic acid (IOP), or depleting DIO2 by shRNA (sh #1, sh #2). **D,** Transwell migration, invasion and 3D invasion assays with A549 cells treated with T3, T4, IOP, T4+I. Results are normalized to Ctrl and shown as mean ± SD (n = 3 for transwell assays and n = 10 for 3D invasion assay). Scale bar (upper two panels), 100µm. Scale bar (bottom panel), 20µm. Statistical methods: two-tailed unpaired t test (transwell); two-tailed Mann-Whitney test (3D invasion). **E,** Transwell migration and invasion assays with SD cells depleting DIO2 by shRNA. Results are normalized to Ctrl and shown as mean ± SD (n = 3). Scale bar, 100µm. Statistical methods: two-tailed unpaired Welch’s t test (migration, Ctrl-sh #1); two-tailed unpaired t test (remaining comparisons). **F,** 3D invasion assay with SD cells treated with IOP or depleting DIO2 by shRNA. Results are normalized to Ctrl and shown as mean ± SD (n = 10). Scale bar, 20µm. Statistical method: two-tailed Mann-Whitney test. **\***, p < 0.05.

### DIO2 and TH enhance the metastatic capabilities of LUAD

DIO2 is a key enzyme involved in metabolism of TH (Fig. 6A). Two major forms of TH, tetraiodothyronine (T4) and triiodothyronine (T3), exist in the human body. DIO2 is mainly responsible for converting T4 to T3, which is much more potent than T4 in activating TH signaling. To provide more evidence that DIO2-TH signaling was the key promoter of EMT and metastasis after phosphorylation, we treated A549 cells with TH to determine whether TH could influence metastatic abilities. As expected, treatment with T3 or T4 elevated M-Markers and attenuated E- Markers in A459 cells, and addition of DIO2 inhibitor IOP disrupted the effect of T4 (Fig. 6B). Besides, treatment with IOP or depleting DIO2 by shRNA in SD cells led to elevation of E- Markers and reduction of M-Markers, suggesting the positive regulation of DIO2 and TH on EMT (Fig. 6C). Next, we performed wound healing, transwell, and 3D invasion assays with A549 cells treated with T3 or T4. We found that T3 and T4 enhanced the metastatic potential of cells without significantly affecting their growth properties (Figs. 6D, S6A-S6B). Notably, the effect of T4 was reversed by IOP, suggesting the dependency of T4 on it. To directly examine the effect of DIO2, we transiently depleted DIO2 in SD cells using siRNA and repeated the same experiments. Depletion of DIO2 mimicked the effect of IOP on inhibiting EMT and invasiveness without significantly affecting cell growth (Figs. S6C-S6E). Finally, stable knockdown of DIO2 by shRNA in SD cells recapitulated the results acquired from transient depletion, confirming the leading role of DIO2 in metastasis (Figs. 6E-6F, S6F-S6G). Taken together, these results demonstrated that DIO2-TH signaling accounted for the metastasis-promoting effect of phosphorylation in LUAD.

### DIO2 and thyropathies are associated with patients’ outcomes

To validate the effect of DIO2 in vivo, we introduced firefly luciferase in SD cells and stably depleted DIO2 (SD-shDIO2 #1 and #2 by two different shRNAs), then intravenously injected SD and SD-shDIO2 cells into nude mice to determine whether depletion of DIO2 or treatment with IOP could attenuate metastasis. As predicted, mice treated with IOP or inoculated with SD-shDIO2 cells consistently displayed fewer metastatic lesions, as shown by the in vivo imaging and staining of lung tissues (Figs. 7A-7B), suggesting suppression of metastasis by targeting DIO2. To seek more clinical support and provide translational values, we obtained LUAD tissue microarray from our cancer center (MCC-TMA) and performed immunohistochemistry staining of DIO2 (Figs. 7C- 7D). We found that H-DIO2 samples (D-Scores ≥ 6) showed a higher rate of metastatic events than L-DIO2 samples (D-Scores < 6), consistent with the results from TCGA-LUAD (Fig. 7E). Interestingly, MCC-TMA also contained 7 patients with hypothyroidism (HypoT) and 1 case with hyperthyroidism (HyperT). Together with the remaining cases (Considered euthyroid, EuT), we compared survival outcomes among three subcohorts. Although not statistically significant, patients in the HypoT group tended to have better survival outcomes than those in the EuT and HyperT groups (Fig. 7F). Due to the scarcity of samples with precise information of thyropathies, this finding was not conclusive, admittedly. Despite this fact, our results provided useful insights into the clinical significance of DIO2 and TH in the metastasis and survival of patients with LUAD.

**Figure 7.**
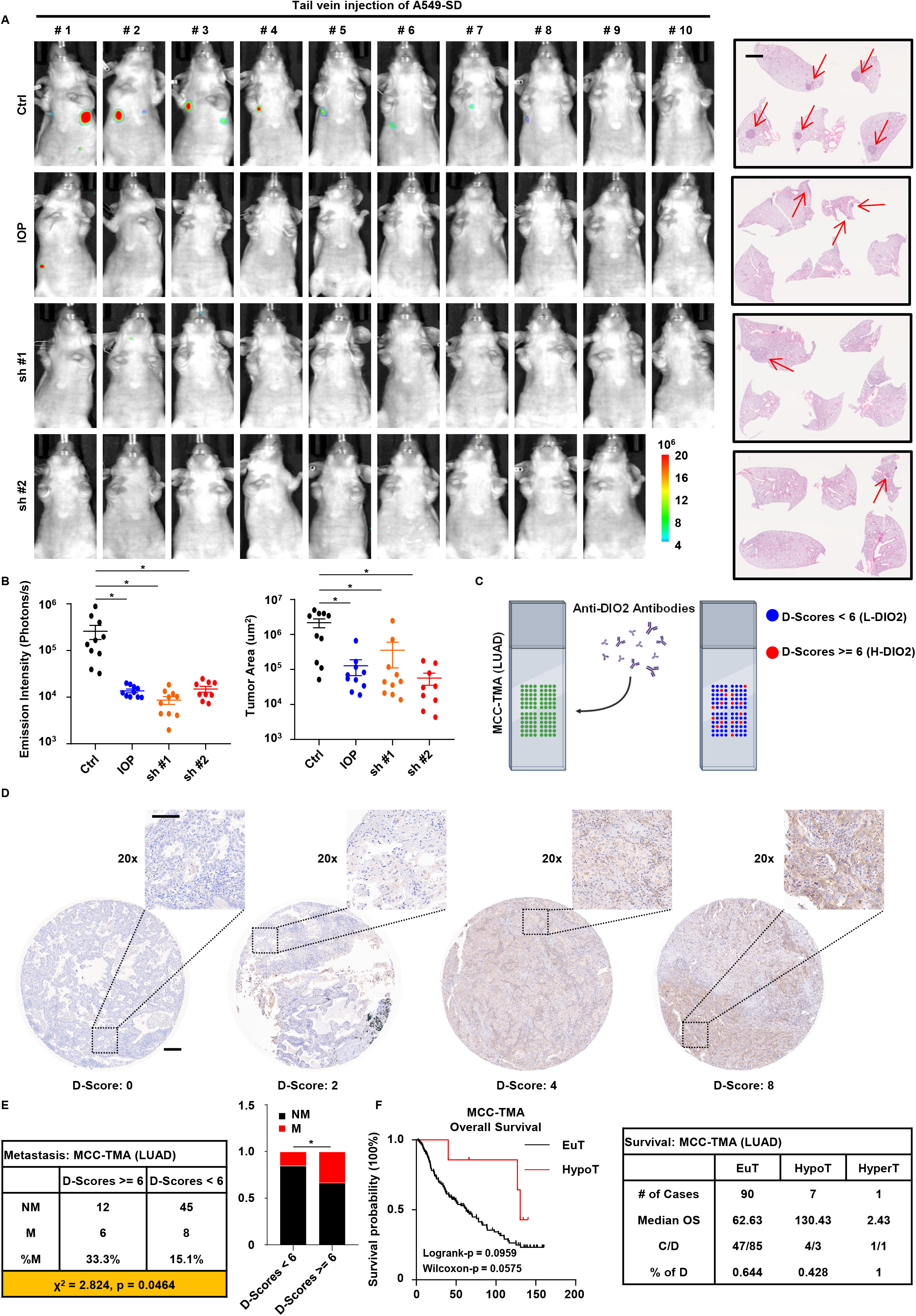
Clinical implication of DIO2 and TH. **A**, (Left panel) In vivo imaging of mice intravenously injected with SD cells depleting DIO2 by shRNA, or mice intravenously injected with SD cells and treated with IOP. (Right panel) H&E staining of lung tissue sections of mice. Scale bar, 2mm. Arrows indicate tumor nodules. **B,** Quantification of results in A. Results are shown as mean ± SEM (n = 10 for Ctrl, IOP, sh #1 and n = 9 for sh #2). Statistical method: two-tailed Mann-Whitney test. **C,** Representation of immunohistochemistry (IHC) staining of DIO2 on Markey Cancer Center Tissue Microarray (MCC-TMA) of LUAD patients, as well as the standard of separating patients based on staining scores of DIO2 (D-Scores). **D,** Representative IHC results of MCC-TMA. Scale bar (origin), 2mm. Scale bar (20x), 100µm. **E,** Analysis of metastatic events of MCC-TMA patients. Based on D- Scores suggested by pathologist, patients are separated into two groups: High DIO2 (D-Scores >= 6, n = 18), Low DIO2 (D-Scores < 6, n = 53). Metastatic events are characterized by regional or distant metastasis. Statistical method: one-sided Chi-square test. **F,** Kaplan-Meier overall survival (OS) curve and statistics of euthyroid (EuT, n = 132) and hypothyroid (HypoT, n = 7) patients in MCC-TMA. The single hyperthyroid (HyperT) case is listed in the statistics table as well. C, censored. D, deaths. Statistical methods: Log-rank test and Gehan-Breslow-Wilcoxon test. **\***, p < 0.05.

## Discussion

Specifically for lung cancer, five-year survival rate for patients with regional diseases is only 30% compared to 60% for patients with localized diseases, and this number further drops to less than 5% for patients with distant metastasis [36]. Sadly, nearly 70% of lung cancer patients develop metastatic lesions upon diagnosis, suggesting an urgent need for more advanced and effective diagnostic techniques for the early stages of metastasis. Here, we provide a possible cause for the metastasis of LUAD (Fig. 8). We show that PLK1 phosphorylates AHR at S489 to promote metastasis. Mechanistically, the Phosphorylation of AHR induces the expression of DIO2, which converts T4 to T3, then activates TH signaling and the EMT process to facilitate the initiation of metastasis. Consistent with the understanding that human and mouse AHR exhibit significant differences in their C-terminal regions [37], the phosphorylation site is likely unique to humans. Currently, PLK1 inhibitors are being tested in phase 2 clinical trials and studies have demonstrated the possibility of targeting PLK1 for cancer treatment with promising efficacy [18, 38–40]. Considering the importance of this phosphorylation event, targeting PLK1 may be an effective treatment for LUAD patients in the future.

**Figure 8.**
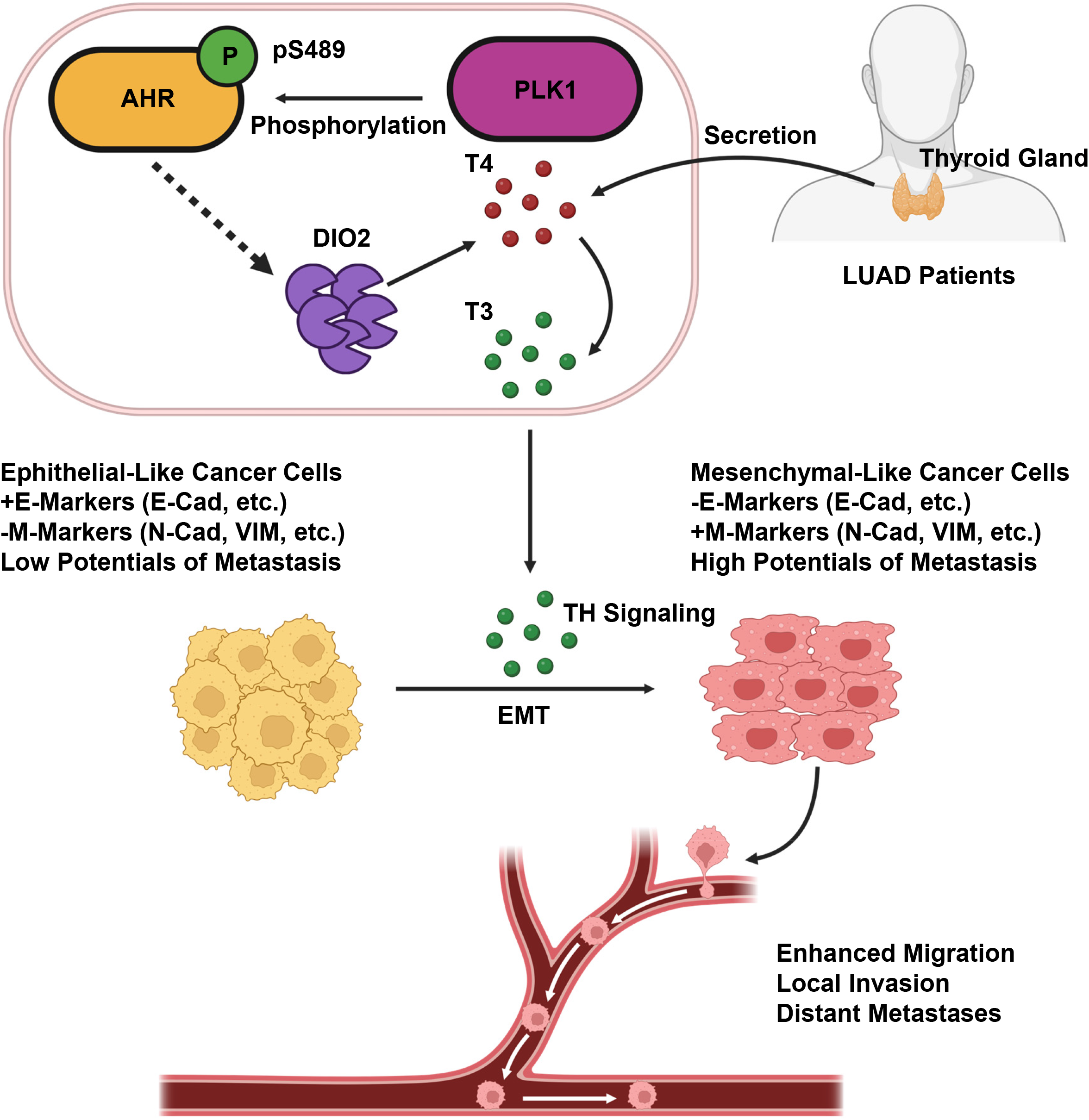
Putative working model. T4 secreted by thyroid glands of LUAD patients enters the circulation system and reaches the local environment of tumors. Phosphorylation of AHR by PLK1 elevates DIO2, which converts the less potent T4 to more potent T3. T3 activates the TH signaling and promotes EMT. Mesenchymal- like tumor cells are prone to local invasion and distant metastasis.

AHR has been shown to account for lung cancer carcinogenesis [41, 42]. However, studies regarding its role in lung cancer metastasis have yielded conflicting conclusions [43]. It is noteworthy that AHR has been shown to have a pro-metastatic function in clinical scenarios, as high levels of AHR are positively associated with disease recurrence and distant metastasis in LUAD patients [23], which is consistent with our results. Another important note is that our working model best fits LUAD and does not support the same notion in LUSC, as the major findings are not repeatable in LUSC. Despite the fact that AHR and PLK1 have been classified as oncogenes in multiple cancers, very few studies have discussed the roles of PLK1 and AHR in LUSC and small cell lung cancer, which comprise 40% of lung cancer cases [44]. Thus, further research is needed to dissect their molecular functions in other subtypes of lung cancer.

Although DIO2 has recently been identified as a metastatic promoter [35], few studies have investigated its function in other cancers. To our knowledge, this study is the first to demonstrate the premetastatic function of DIO2 in LUAD. One question that remains unanswered is how phosphorylation increases the level of DIO2. As AHR is a transcription factor, it is rational to hypothesize that phosphorylation may elevate the activity of AHR and enhance the transcription of DIO2. However, this is unlikely to happen in our model, as neither phosphorylation elevates the transcriptional activity of AHR nor treatment with an AHR inhibitor decreases the level of DIO2 (data not shown). Notably, AHR has extensive crosstalk with other pathways and can serve as a cofactor for several other transcription factors [45, 46]. Thus, it is possible that phosphorylated AHR cooperates with these transcription factors to jointly elevate the level of DIO2, and this possibility remains to be explored.

Our results support the idea that hypothyroidism may be a good prognostic factor in lung cancer patients, consistent with a previous report that NSCLC patients with hypothyroidism have better overall survival than euthyroid patients [47]. In fact, as one of the complications frequently induced by immune checkpoint inhibitors, patients with hypothyroidism respond better to anti- PD-1 immunotherapy, supporting the notion that hypothyroidism is a strong indicator of better outcomes [48]. Interestingly, targeting both PLK1 and AHR have been evaluated as strategies to enhance the effect of immune checkpoint inhibitors [49, 50]. Thus, it is possible that phosphorylation modulates the level of TH in patients and leads to an unfavorable response to checkpoint inhibitors. Considering this possibility, future studies should specifically address this question and provide suggestions for developing novel clinic-based treatments targeting these important proteins.

## Acknowledgement

This study is supported by NIH R01 CA157429 (XL), R01 CA196634 (XL), R01 CA264652 (XL), R01 CA256893 (XL), as well as the Biospecimen Procurement & Translational Pathology, Biostatistics and Bioinformatics, Redox Metabolism, and Flow Cytometry and Immune Monitoring Shared Resources of the University of Kentucky Markey Cancer Center (P30CA177558).

## Data availability

All data are included in this article and the original data are available from the corresponding author upon reasonable request.

---

**Figure S1.**
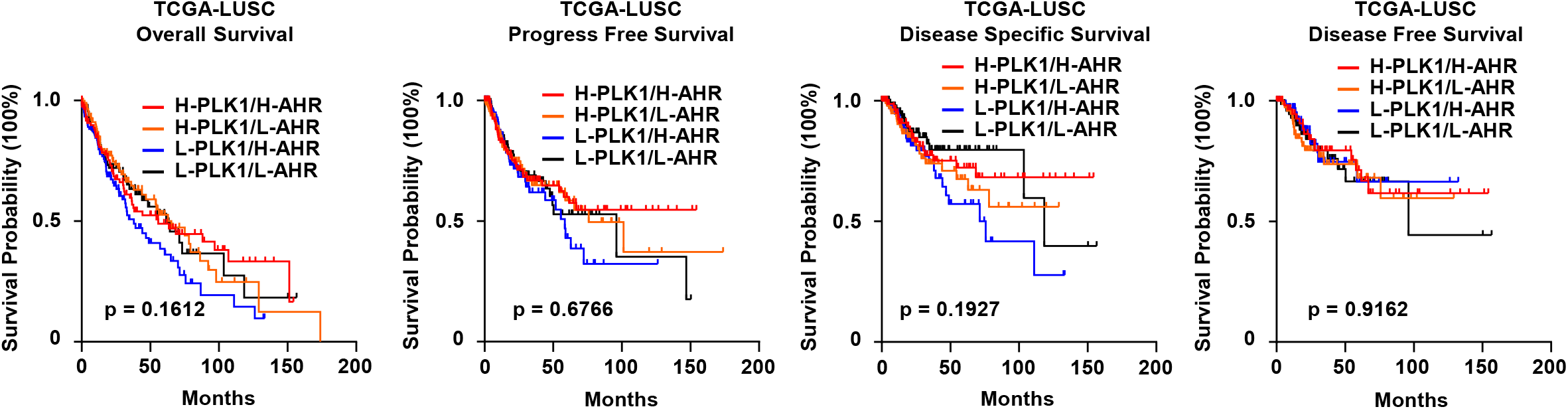
Kaplan-Meier survival curves of TCGA-LUSC patients. Based on the median expressions of PLK1 and AHR, patients are separated into four groups: High PLK1/High AHR (H-PLK1/H-AHR, PLK1/AHR > median, n = 118/118/102/61 from left to right), High PLK1/Low AHR (H-PLK1/L-AHR, PLK1 > median/AHR < median, n = 122/122/109/78 from left to right), Low PLK1/High AHR (L-PLK1/H-AHR, PLK1 < median/AHR > median, n = 122/123/110/72 from left to right), Low PLK1/Low AHR (L-PLK1/L-AHR, PLK1/AHR < median, n = 116/116/108/81 from left to right). Statistical method: Log-rank test.

**Figure S2.**
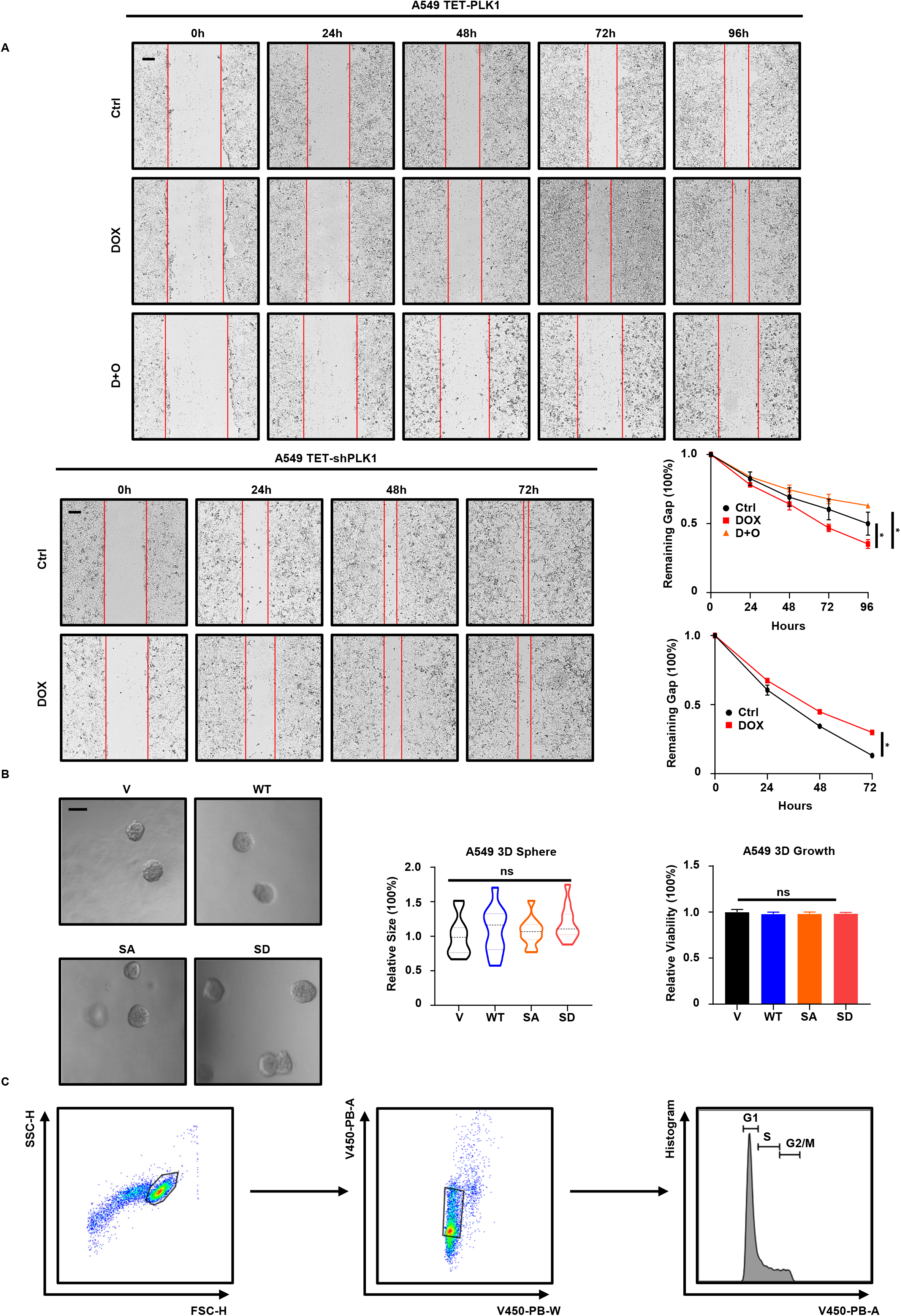
PLK1 phosphorylates AHR to promote metastasis of LUAD. **A**, Wound healing assays with A549 cells with TET-PLK1 or TET-shPLK1. D+O, doxycycline plus onvansertib. Results are normalized to 0h and shown as mean ± SD (n = 3). Scale bar, 250µm. Statistical method: two-tailed unpaired t test. **B,** 3D spheroid formation and 3D growth assays with V, WT, SA, SD cells. Results are normalized to V and shown as mean ± SD (n = 11 for spheroid formation and n = 6 for 3D growth). Scale bar, 20µm. Statistical method: one-way ANOVA test. **C,** Gating strategy of cell cycle analysis by flow cytometry. 10^4^ cells are counted for each group. **\***, p < 0.05. ns, not significant.

**Figure S3.**
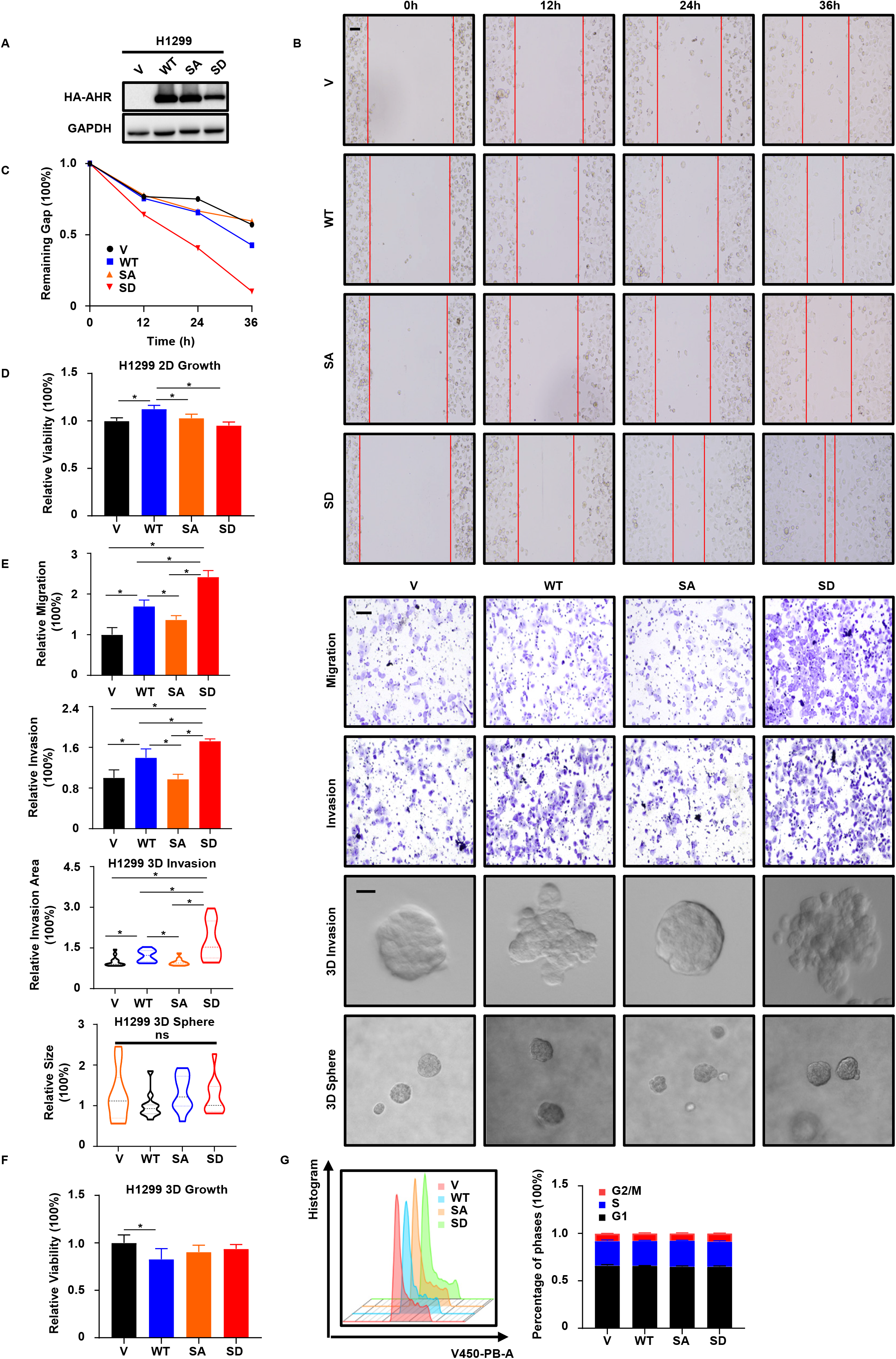
Metastasis assays with H1299 cells. **A**, IB to verify establishment of V, WT, SA, SD cells. **B, C,** Wound healing assay with V, WT, SA, SD cells. Results are normalized to 0h. Scale bar, 100µm. **D,** 3-day 2D growth assay with V, WT, SA, SD cells. Results are normalized to day 0 and shown as mean ± SD (n = 8). Statistical method: one-way ANOVA test following multiple comparisons. **E,** Transwell migration, invasion, 3D invasion and spheroid formation assays with V, WT, SA SD cells. Results are normalized to V and shown as mean ± SD (n = 3 for transwell, n = 12 for 3D invasion, n = 11 for spheroid formation). Scale bar (upper two panels), 100µm. Scale bar (bottom two panel), 20µm. Statistical methods: two-tailed unpaired t test (transwell); two-tailed Mann-Whitney test (3D invasion); Kruskal-Wallis test following multiple comparisons (3D sphere). **F,** 3D growth assay with V, WT, SA, SD cells. Results are normalized to V and shown as mean ± SD (n = 6). Statistical method: one-way ANOVA test following multiple comparisons. **G,** Cell cycle analysis of V, WT, SA, SD cells by flow cytometry. Results are normalized to V and shown as mean ± SD (n = 3). **\***, p < 0.05.

**Figure S4.**
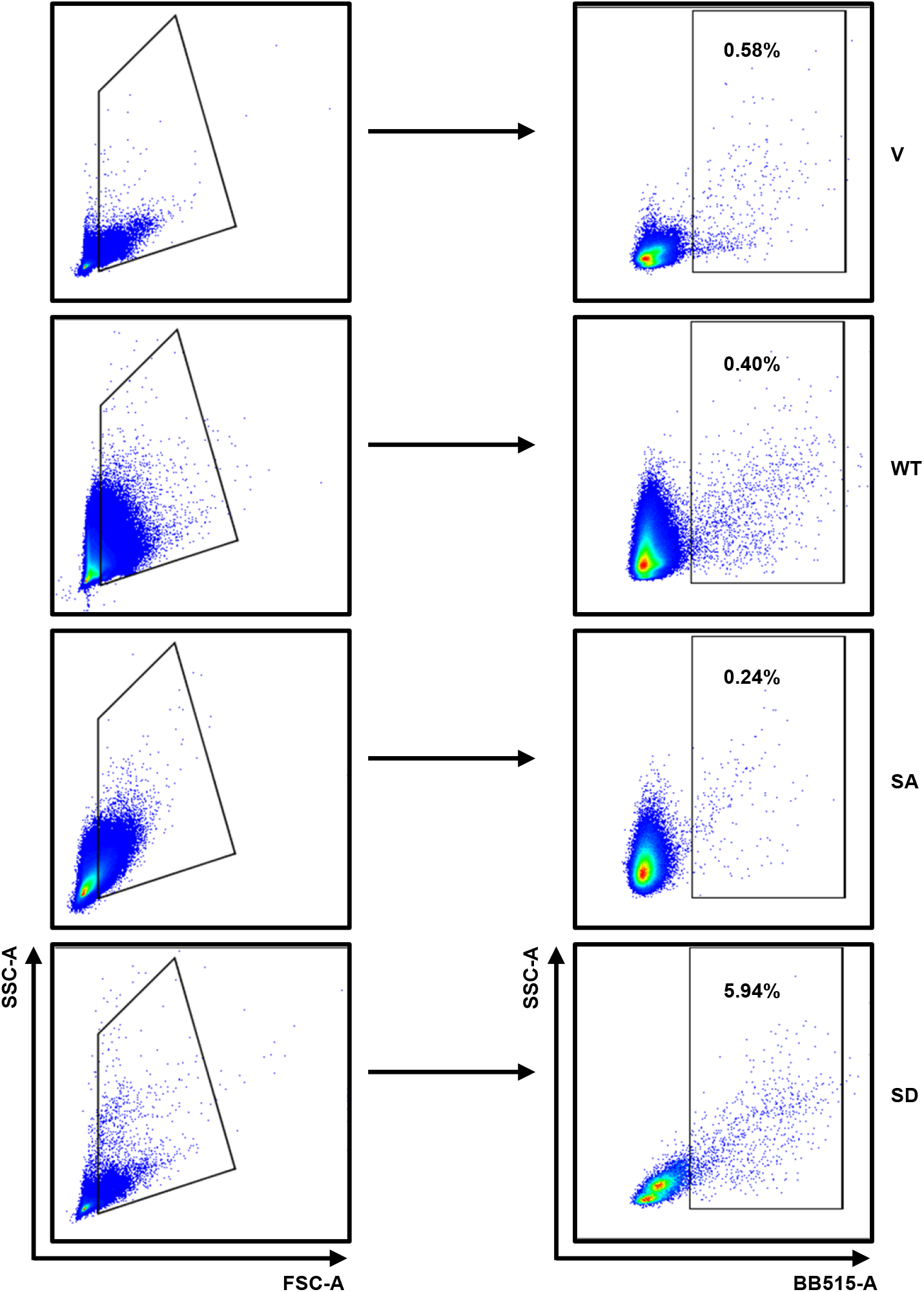
Gating strategy of flow cytometry analysis of circulating GFP+ tumor cells. Each sample is enriched with 10^3^ GFP^+^ cells and the percentage of GFP^+^ tumor cells is calculated.

**Figure S5.**
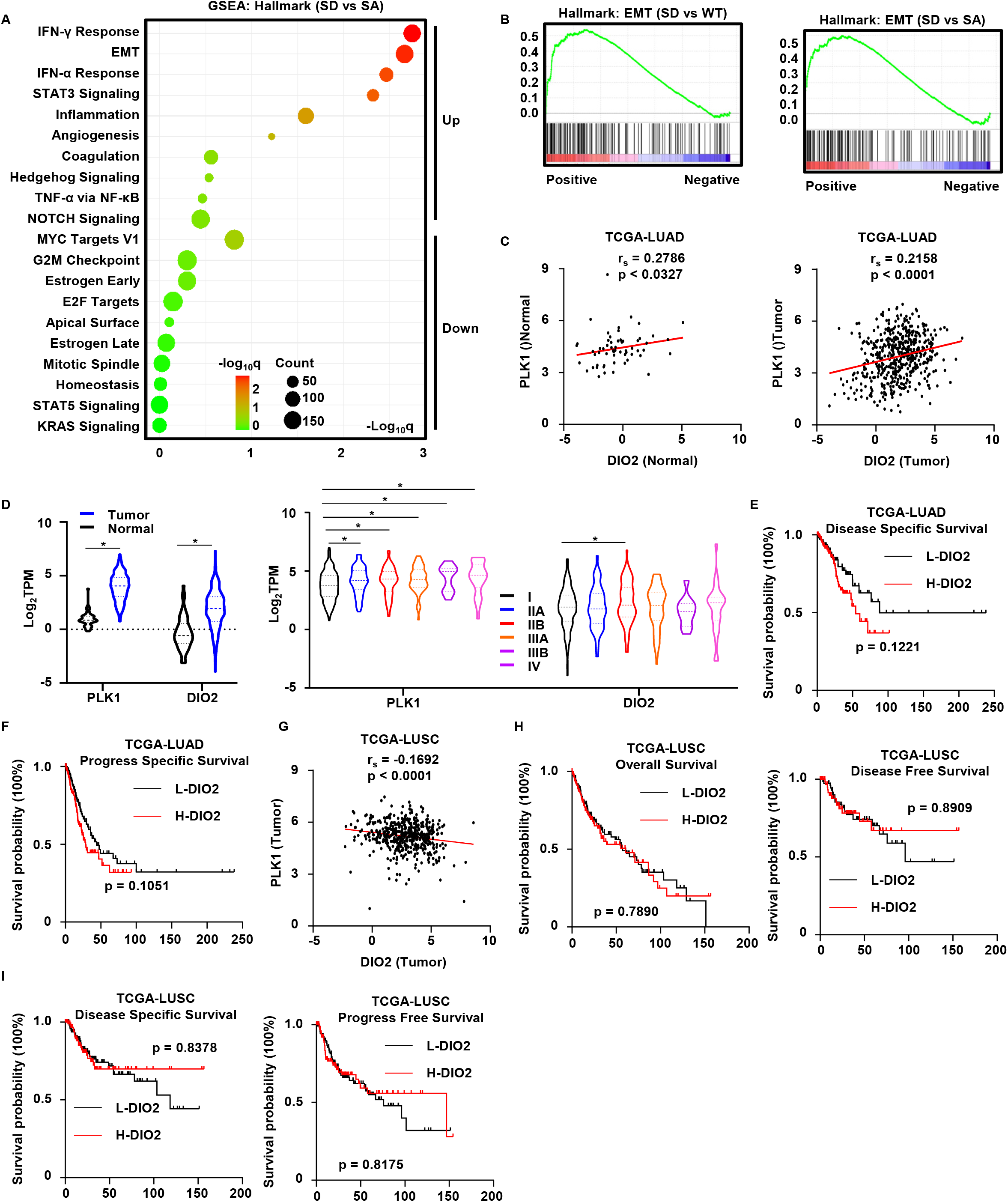
RNA-seq analysis and validation. **A**, Bubble plot (SD vs SA) of top upregulated and downregulated pathways in SD cells, identified by GSEA with hallmark gene set. **B,** Enrichment plots of EMT pathway. **C,** Spearman correlation analysis of DIO2 and PLK1 in normal lung samples (n = 59) and tumor samples (n = 513) from TCGA-LUAD. r_s_, spearman correlation coefficient. **D,** Expressions of DIO2 and PLK1 between normal lung samples and tumor samples, as well as among different stages of tumor samples, from TCGA-LUAD. Log_2_TPM, log_2_-transformed Transcripts Per Million values. Statistical methods: linear-mixed model test (left panel); one-tailed Mann-Whitney test (right panel). **E, F,** Kaplan-Meier survival curves of TCGA-LUAD patients. H-DIO2, n = 118 (E)/125 (F). L-DIO2, n = 112 (E)/125 (F). Statistical method: Log-rank test. **G,** Spearman correlation analysis of DIO2 and PLK1 in tumor samples (n = 501) from TCGA-LUSC. **H, I,** Kaplan-Meier survival curves of TCGA-LUSC patients. Based on the 1^st^ and 3^rd^ quartile expression of DIO2, patients are separated into two groups: High DIO2 (H-DIO2, >= 3^rd^ quartile) and Low DIO2 (L-DIO2, <= 1^st^ quartile). H-DIO2, n = 120 (H, left panel)/67 (H, right panel)/107 (I, left panel)/120 (I, right panel). L-DIO2, n = 120 (H, left panel)/77 (H, right panel)/109 (I, left panel)/120 (I, right panel). Statistical method: Log-rank test. **\***, p < 0.05.

**Figure S6.**
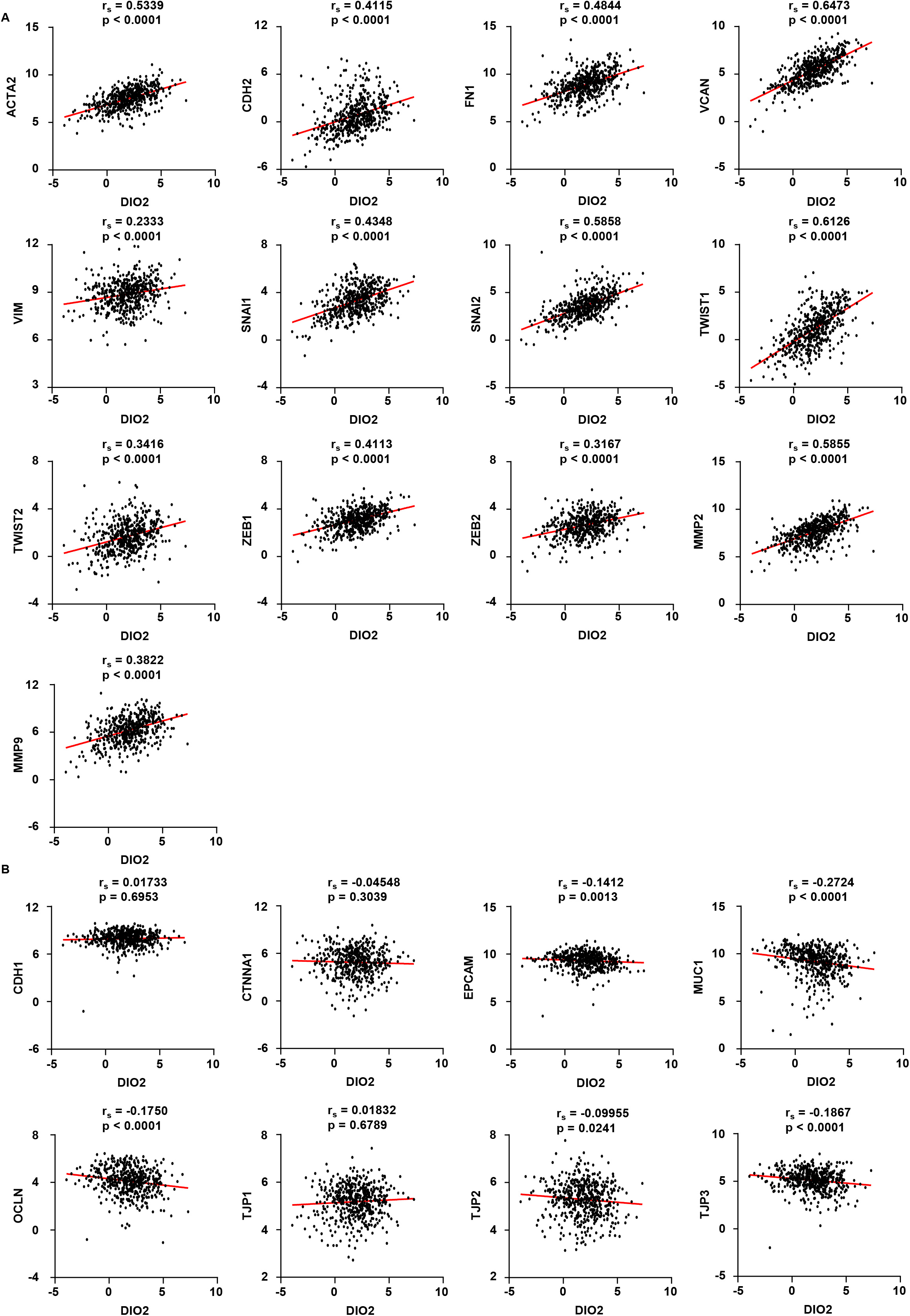
Spearman correlation analysis of DIO2 and EMT markers in TCGA-LUAD. **A**, Correlation analysis of DIO2 and M-Markers. **B,** Correlation analysis of DIO2 and E-Markers.

**Figure S7.**
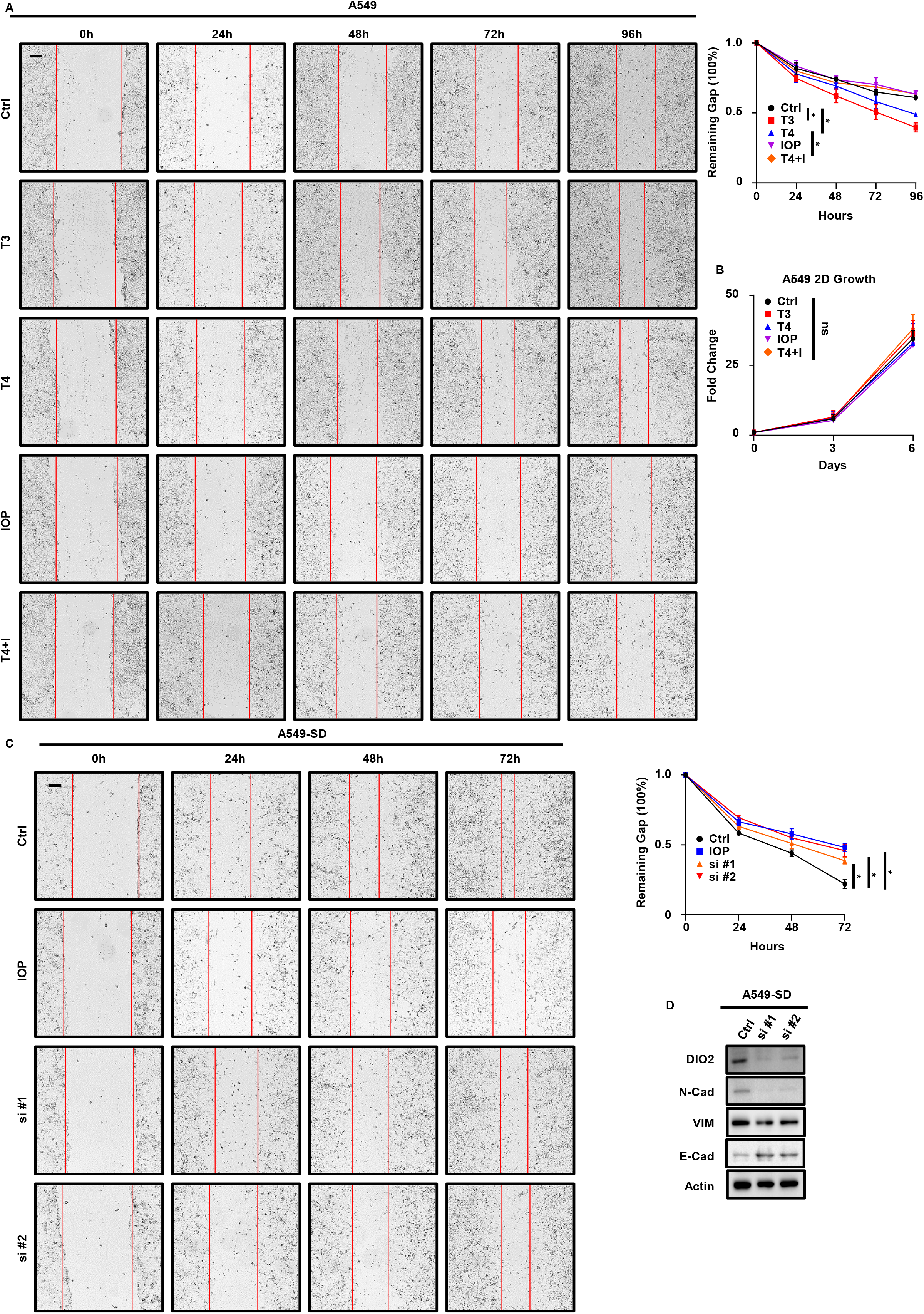
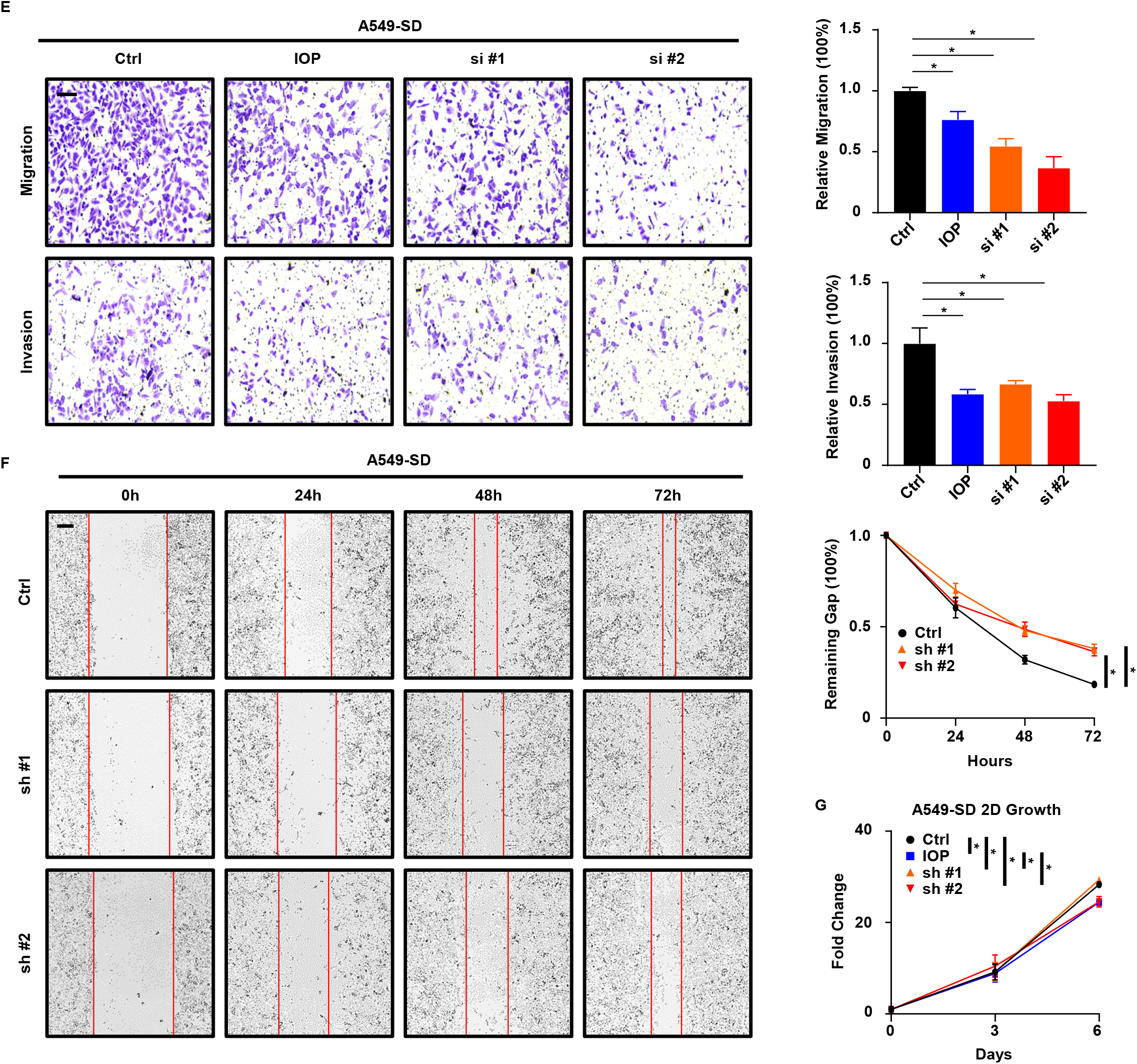
DIO2 promotes metastasis of LUAD via TH signaling. **A**, Wound healing assay with A549 cells treated with T3, T4, IOP, T4+I. Results are normalized to 0h and shown as mean ± SD (n = 3). Scale bar, 250µm. Statistical method: two-tailed unpaired Welch’s t test (T3-V); two-tailed unpaired t test (T4-V, T4+I-T4). **B,** 2D cell growth assay with A549 cells treated with T3, T4, IOP, T4+I. Results are normalized to day 0 and shown as mean ± SD (n = 8). Statistical method: Kruskal-Wallis test. **C,** Wound healing assay with SD cells treated with IOP or transiently depleting DIO2 by siRNA (si #1, si #2). Results are normalized to 0h and shown as mean ± SD (n = 3). Scale bar, 250µm. Statistical method: two-tailed unpaired t-test. **D,** IB to detect EMT markers in SD cells transiently depleting DIO2 by siRNA. **E,** Transwell migration and invasion assays with SD cells treated with IOP or depleting DIO2 by siRNA. Results are normalized to Ctrl and shown as mean ± SD (n = 3). Scale bar, 100µm. Statistical method: two-tailed unpaired t test. **F,** Wound healing assay with SD cells depleting DIO2 by shRNA. Results are normalized to 0h and shown as mean ± SD (n = 3). Scale bar, 250µm. Statistical method: two-tailed unpaired t test. **G,** 2D cell growth assay with A549 cells treated with IOP or depleting DIO2 by shRNA. Results are normalized to day 0 and shown as mean ± SD (n = 8). Statistical method: Welch’s ANOVA test following multiple comparisons. **\***, p < 0.05. ns, not significant.

**Table S1.** Survival information of TCGA-LUAD patients.

| Table S1. Survival information of TCGA-LUAD patients |  |  |  |  |
| --- | --- | --- | --- | --- |
|  | TCGA-LUAD |  |  |  |
|  | H-PLK1/H-AHR | H-PLK1/L-AHR | L-PLK1/H-AHR | L-PLK1/L-AHR |
| Overall Survival |  |  |  |  |
| Median Survival | 36.66 | 47.80 | 56.71 | 53.65 |
| Censored/Events | 51/48 | 87/59 | 110/37 | 69/36 |
| % of Events | 0.485 | 0.404 | 0.252 | 0.343 |
| Significance | Yes |  |  |  |
| Progress Free Survival |  |  |  |  |
| Median Survival | 23.84 | 37.61 | 44.05 | 36.13 |
| Censored/Events | 48/51 | 89/57 | 96/51 | 61/44 |
| % of Events | 0.515 | 0.390 | 0.347 | 0.419 |
| Significance | Yes |  |  |  |
| Disease Specific Survival |  |  |  |  |
| Median Survival | 40.41 | N/A (>50%) | N/A (>50%) | 88.14 |
| Censored/Events | 60/32 | 97/35 | 117/25 | 78/18 |
| % of Events | 0.348 | 0.265 | 0.176 | 0.188 |
| Significance | Yes |  |  |  |
| Disease Free Survival |  |  |  |  |
| Median Survival | 35.61 | 96.95 | N/A (>50%) | 158.20 |
| Censored/Events | 29/24 | 60/20 | 76/25 | 44/18 |
| % of Events | 0.453 | 0.250 | 0.248 | 0.290 |
| Significance | Yes |  |  |  |

**Table S2.** Survival information of TCGA-LUSC patients.

| Table S2. Survival information of TCGA-LUSC patients |  |  |  |  |  |
| --- | --- | --- | --- | --- | --- |
|  |  | TCGA-LUSC |  |  |  |
|  |  | H-PLK1/H-AHR | H-PLK1/L-AHR | L-PLK1/H-AHR | L-PLK1/L-AHR |
| Overall Survival |  |  |  |  |  |
|  | Median Survival | N/A (>50%) | 75.78 | 58.49 | 96.10 |
|  | Censored/Events | 84/33 | 86/36 | 85/38 | 84/32 |
|  | % of Events | 0.282 | 0.295 | 0.309 | 0.276 |
|  | Significance | No |  |  |  |
| Progress Free Survival |  |  |  |  |  |
|  | Median Survival | N/A (>50%) | N/A (>50%) | 75.75 | 118.36 |
|  | Censored/Events | 83/18 | 86/23 | 84/26 | 93/15 |
|  | % of Events | 0.178 | 0.211 | 0.236 | 0.139 |
|  | Significance | No |  |  |  |
| Disease Specific Survival |  |  |  |  |  |
|  | Median Survival | N/A (>50%) | N/A (>50%) | N/A (>50%) | 96.10 |
|  | Censored/Events | 47/14 | 60/18 | 60/12 | 65/16 |
|  | % of Events | 0.230 | 0.231 | 0.167 | 0.198 |
|  | Significance | No |  |  |  |
| Disease Free Survival |  |  |  |  |  |
|  | Median Survival | 55.73 | 65.23 | 37.94 | 61.61 |
|  | Censored/Events | 66/51 | 73/49 | 62/60 | 73/43 |
|  | % of Events | 0.436 | 0.402 | 0.492 | 0.371 |
|  | Significance | No |  |  |  |

**Table S3.** Survival information of patients with H-DIO2 and L-DIO2.

| Table S3. Survival information of patients with H-DIO2 and L-DIO2 |  |  |  |  |
| --- | --- | --- | --- | --- |
|  | TCGA-LUAD |  | TCGA-LUSC |  |
|  | H-DIO2 | L-DIO2 | H-DIO2 | L-DIO2 |
| **Overall Survival** |  |  |  |  |
| Median Survival | 42.34 | 59.11 | 60.52 | 57.07 |
| Censored/Events | 75/50 | 83/40 | 70/50 | 64/56 |
| % of Events | 0.4 | 0.325 | 0.471 | 0.467 |
| Significance | Yes |  | No |  |
| **Progress Free Survival** |  |  |  |  |
| Median Survival | 28.44 | 44.05 | 146.99 | 75.78 |
| Censored/Events | 71/54 | 77/46 | 87/33 | 80/40 |
| % of Events | 0.432 | 0.346 | 0.275 | 0.333 |
| Significance | No |  | No |  |
| **Disease Specific Survival** |  |  |  |  |
| Median Survival | 54.34 | 88.14 | N/A (>50%) | 118.355 |
| Censored/Events | 85/33 | 87/23 | 87/20 | 83/26 |
| % of Events | 0.28 | 0.209 | 0.187 | 0.238 |
| Significance | No |  | No |  |
| **Disease Free Survival** |  |  |  |  |
| Median Survival | 62.23 | N/A (>50%) | N/A (>50%) | 96.1 |
| Censored/Events | 44/23 | 53/16 | 54/13 | 58/19 |
| % of Events | 0.343 | 0.232 | 0.194 | 0.246 |
| Significance | Yes |  | No |  |

## Notes

https://www.cbioportal.org/

https://portal.gdc.cancer.gov/projects/TCGA-LUAD

### Competing Interest Statement

The authors have declared no competing interest.

